# Traditional medicinal use is linked with apparency, not specialized metabolite profiles in the order Caryophyllales

**DOI:** 10.1101/2023.07.22.550123

**Authors:** Alex H. Crum, Lisa Philander, Lucas Busta, Ya Yang

**Author notes:** Author for correspondence: Alex H. Crum.

## Abstract

A better understanding of the relationship between plant specialized metabolism and traditional medicinal use has the potential to aid in bioprospecting and the untangling of cross-cultural plant use patterns. However, given the limited information available for metabolites in most plant species, associating medicinal properties with a metabolite can be difficult. The order Caryophyllales has a unique pattern of lineages of tyrosine- or phenylalanine-dominant specialized metabolism, represented by mutually exclusive anthocyanin and betalain pigments, making the group ideal to work around a lack of detailed knowledge of specific metabolites. We compiled a list of medicinal species in selected tyrosine- or phenylalanine-dominant families of Caryophyllales across the globe (Nepenthaceae, Polygonaceae, Simmondsiaceae, Microteaceae, Caryophyllaceae, Amaranthaceae, Limeaceae, Molluginaceae, Portulacaceae, Cactaceae, and Nyctaginaceae) by searching scientific literature until no new uses were recovered, and tested for phylogenetic clustering of medicinal uses using a “hot nodes” approach. To test potential non-metabolite drivers of medicinal use, like how often humans encounter a species (apparency), we repeated the same analysis in North American species across the entire order and performed phylogenetic generalized least squares regression (PGLS) with occurrence data from the Global Biodiversity Information Facility (GBIF). We hypothesized families with Tyr-enriched metabolism would show clustering of different types of medicinal use compared to the ancestral Phe-enriched metabolism. Instead, weedy, wide-ranging clades in Polygonaceae and Amaranthaceae are overrepresented across nearly all types of medicinal use. Therefore, we found that apparency is a better predictor of medicinal use than metabolite profiles, although metabolism type may still be a contributing factor.

## INTRODUCTION

Nearly 80% of the population in low and middle income countries relies on medicinal plants as their primary form of healthcare (Gaoue et al., 2021; Hamilton, 2004). Globally, even those not directly relying on medicinal plants will access pharmaceuticals developed from plant-based natural product extractions. Natural products in pharmacological studies are also known as specialized metabolites (SMs), reflecting the chemical’s function in the plant rather than their service to people. SMs are important for plant survival, helping mediate biotic and abiotic stress responses and playing a role in plant communications (e.g. attracting pollinators).

These compounds are extremely diverse, and it is estimated that the plant kingdom produces over one million unique SMs (Afendi et al., 2012). Specialized metabolites tend to be present in characteristic phylogenetic patterns – while some are relatively widespread, such as anthocyanins, others are more lineage specific, such as the glucosinolates mostly restricted to Brassicales. Additionally, closely related plants tend to produce similar SMs, being constrained by the SM biosynthetic pathways of their common ancestor (Wink, 2003).

Species used in traditional medicine are also clustered into certain phylogenetic lineages, with a species tending to be related to plants with similar use more often than random chance (Gaoue et al., 2021; Saslis-Lagoudakis et al., 2011, 2012). Because a plant’s medicinal effect is determined by its underlying specialized metabolism and closely related plants tend to produce similar SMs, this phylogenetic pattern of traditional use likely reflects phylogenetic clustering of medicinal SMs (Pellicer et al., 2018). Therefore, traditional medicinal use in a clade should reflect the classes of SMs to be found in a clade, potentially guiding bioprospecting for new drugs to treat a variety of conditions. Reciprocally, knowing the specialized metabolism in a clade has the potential to allow ethnobotanists to predict and prioritize investigation of medicinal species in a given geographical region.

However, associating a particular SM or class of SMs with a particular medicinal use has proven to be a challenging, time-consuming process (David et al., 2015). There are gaps in reported knowledge in both ethnobotanical uses and metabolomics/natural products studies.

Additionally, the correlation between a SM and medicinal use is likely not a linear one. Potential interactions of multiple plants in a medicinal recipe or the use of one plant to boost another’s effect cannot be discounted, as this could contribute to a medicinal effect in a way individual species would not and may not be reported in literature. For example, interactions between *Psychotria viridis* and *Banisteriopsis caapi* in a common Ayahuasca recipe are what give it its psychoactive effect. The active compound from *P. viridis* would not be able to reach the central nervous system without SMs from *B. caapi* protecting it from degradation, a relationship that has only been understood after extensive investigation of these plants’ SMs (Gambelunghe et al., 2008). It is also likely that not all of the plant species in the indigenous pharmacopeia have a pharmacological effect matching their reported uses.

Although most of the challenges of associating SM and traditional medicinal use across a phylogeny cannot be dealt with in a single study, the order Caryophyllales is a compelling system to work around the lack of known SMs in most plants due to its unique evolution of tyrosine- and phenylalanine-dominant specialized metabolism. Consisting of approximately 12,500 species across 39 families, Caryophyllales includes many well-known species used in traditional medicine, such as jojoba (*Simmondsia chinensis; Simmondsiaceae*), peyote (*Lophophora williamsii*; Cactaceae), and Dock (*Rumex* spp.; Polygonaceae). Although suffering from the same lack of known metabolites as all other plant groups, the active constituents of some of the order’s more charismatic plants have been identified. For example, mescaline is known to be the SM that confers peyote’s hallucinogenic effect, while epinephrine and morphine are both well-known human drugs that are synthesized from tyrosine (Tyr) in plants. Mescaline, morphine, and epinephrine are also emblematic of a unique evolution of specialized metabolism in the order, which is also responsible for the Caryophyllales-specific betalain pigments.

Both mescaline and the betalains, which range from yellow to pink, are derived from the amino acid Tyr, and are part of a Tyr-derived SM diversification in the order. This diversification is the result of the duplication of arogenate dehydrogenase (ADH), the gene responsible for converting Tyr from its precursor, arogenate, at the base of the core Caryophyllales clade (Lopez-Nieves et al., 2018). While the canonical version (ADH-β) is feedback-inhibited by excess tyrosine, the new copy (ADH-α) has relaxed that regulation. Species with ADH-α therefore have an excess of tyrosine, which the core Caryophyllales has incorporated into specialized metabolism in a variety of ways, including isoquinoline alkaloids such as mescaline, betalains, and catecholamines (Lopez-Nieves et al., 2018). These SMs, in turn, help plants mediate both abiotic and biotic stressors such as UV light and herbivory. At least in the case of the betalains, there is some evidence that these SMs have some advantages over their non-tyrosine derived functional homologues in other plants, such as being more pH stable, and therefore provides more protection for CAM plants than the more common anthocyanin pigments (Jain and Gould, 2015).

Conversely, arogenate is also the precursor to the amino acid phenylalanine (Phe), which is itself a major precursor to phenylpropanoid SMs, such as flavonoids, lignins, and anthocyanins. These SMs are canonically a huge component of specialized metabolism overall, aiding in structure, pollinator attraction, and defense. For example, anthocyanin pigments, which range from red to blue and in contrast to betalains are widespread across the plant kingdom, play many roles, including photoprotection, pollinator attraction, and antioxidant action (Landis et al., 2015). However, because Tyr and Phe are competing for the same precursor, the duplication and relaxed regulation of ADH in the Caryophyllales core clade means that there is less arogenate being fed into the Phe synthesis and the downstream pathways.

One apparent consequence of this competition between Tyr- and Phe-dominant SM pathways is the mutual exclusivity of betalains and anthocyanins in the order. Betalains, derived from Tyr, are restricted to the core Caryophyllales clade. However, the anthocyanins, derived from Phe, are ubiquitous throughout the plant kingdom. In lineages that have betalain, anthocyanin is never found, and vice versa (Clement and Mabry, 1996). Although the gain of ADH-α appears to have occurred only once, betalain itself likely had multiple origins in the core clade, leading to a pattern of anthocyanin-producing lineages sister to betalain producing ones (Sheehan et al., 2020 citation; **Fig. 1).** These anthocyanin-producing lineages in the core clade (the most notable being the order’s namesake, Caryophyllaceae) tend to eventually lose the function of their ADH-α or lose it entirely, making the type of pigment a species produces a good proxy for presence/absence of ADH-α and therefore also a shorthand for overall metabolism type (Phe- vs Tyr-dominant) (Lopez-Nieves et al., 2018).

**Fig. 1.**
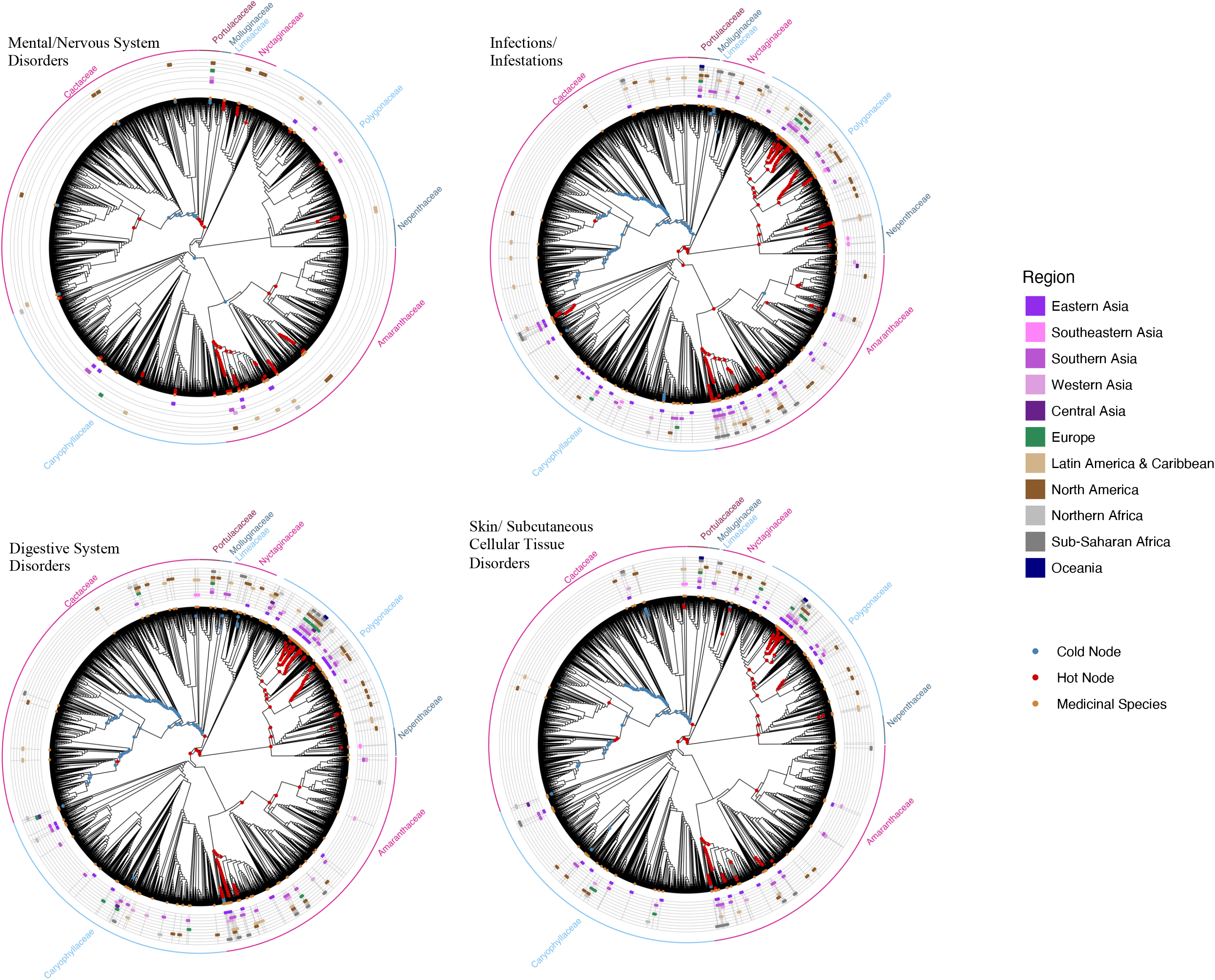
A family level phylogeny of Caryophyllales. Blue branches represent anthocyanin producing lineages, while magenta represents betalain producing ones. Families surveyed in the global medicinal use literature review are outlined in black. Representatives of well known medicinal species are shown next to the phylogeny. PCs: Bruce Ackley, Ohio State University, Bugwood.org; Jeff Abbas 2006; Monty Rickard 2003.

This unusual tension between SM pathways makes Caryophyllales an ideal system to study the relationship between specialized metabolism and human medicinal use. We predicted that within Caryophyllales, lineages with different dominant metabolic pathways would be associated with different categories of medicinal uses. For example several Tyr-derived SMs such as salidroside, mescaline, and catecholamines affect the nervous system, while there is some evidence supporting anti-cancer, cardiovascular, and dermatological benefits of Phe-derived SMs such as the stilbenoid resveratrol and flavonoid phloretin (Berman et al., 2017; Casarini et al., 2020; Vamvakopoulou et al., 2023, Zhong et al., 2018).

To test the hypothesis that lineages with Tyr- and Phe-dominant specialized metabolism have different categories of use in traditional medicine, we used a phylogenetic hot nodes approach to explore over- and under-representation of different categories of medicinal use in selected lineages of the order, with representatives from both betalain and anthocyanin-producing families (Pellicer et al., 2018; Saslis-Lagoudakis et al., 2012; Zaman et al., 2021). We found that contrary to predictions, rather than different categories being overrepresented in betalain clades vs. the anthocyanin clades, the same clades are overrepresented in nearly every category. To test whether a factor other than metabolism, such as proximity to humans, may be driving medicinal plant selection, we used Global Biodiversity Information Facility (GBIF) occurrence data on all North American members of the Caryophyllales to explore the relationship between medicinal use and how apparent (frequently encountered and noticed) a species is to humans via phylogenetic generalized least squares regression (PGLS).

## METHODS

### Literature Search

Caryophyllales accounts for roughly 6% of all known angiosperm species (Thulin et al., 2016). Given the size of the order, assembling all known medicinal uses is prohibitive. Therefore, we selected families that represent the phylogenetic and metabolic diversity of the Caryophyllales (Fig. 1). These include ancestral anthocyanin-producing families (Polygonaceae, Nepenthaceae, and Simmondsiaceae), betalain-producing families (Cactaceae, Portulacaeae, and Amaranthaceae), and anthocyanin-producing families in core Caryophyllales representing reversals to Phe-enriched metabolism (Caryophyllaceae, Limeaceae, Molluginaceae, and Microteaceae). Apart from representing either Phe- or Tyr-dominant specialized metabolism using contrasting pigment types as a proxy, these families either have well-known species with medicinal use (i.e. San Pedro cactus, rhubarb, and chickweed), or are of manageable sizes. Additionally, to mitigate the potential of a specific climate selecting for metabolites of a given effect, these families represent a range of climatic niches, with desert, tropical, and temperate species represented in selected anthocyanin and betalain producing families.

A review of the medicinal uses in these families across the globe was conducted using Google Scholar, the Native American Ethnobotany Database (NAEB), and the PubMed database via the R package rentrez (Moerman, 2003; Winter, 2017). When taxonomy in the reported medicinal use differed from species name on the guide tree (see methods below), it was checked using Plants of the World and Tropicos and reconciled (POWO, 2023; Tropicos). Specific medicinal use and culture/region were recorded for each species. Medicinal uses were then categorized into broader categories based on physiological system (e.g. respiratory, cardiovascular, or digestive) according to Level 2 of the Economic Botany Data Collection Standard (Cook, 1995). The region of reported use for a plant species was categorized using the M49 standard of the United Nations Statistics Division (Supplementary Table 1).

### Testing for phylogenetic clustering

Phylogenetic clustering of medicinal use was tested using a “hot nodes” approach (Ernst et al., 2016; Saslis-Lagoudakis et al., 2012). In this method, nodes on a phylogeny with statistical overrepresentation of species with a reported medicinal use in a given category are “hot”. Additionally, nodes with statistical underrepresentation are “cold” nodes. The Caryophyllales phylogeny as published in Smith et al. (2018) was trimmed to the selected families using Phyx to serve as the guide tree for analyses (Brown et al., 2017). Over- and under-representation of each category of medicinal use across this guide tree was tested using the “nodesig” command in Phylocom v4.2 (Webb et al., 2008). Phylocom output, region of use, and reported species were visualized on the phylogeny in R using the ggtree and ggtreeExtra packages (Yu et al., 2017; Xu et al., 2021).

### Phylogenetic regression of North American medicinal use

To explore the relationship between medicinal use and apparency to humans, medicinal uses exclusively from North America were collected and tested for hot nodes as described above for the entirety of Caryophyllales. Additionally, occurrence records of all Caryophyllales species from North America were downloaded from GBIF and cleaned and quality checked using the R package CoordinateCleaner (Zizka et al., 2019). Occurrences belonging to species not on the guide tree were discarded. Reciprocally, species not in the occurrence dataset were trimmed from the guide tree. Species on the tree were classified as “medicinal” or “not medicinal”. Occurrence records for each species were counted and used directly in PGLS as a proxy for apparency, which was tested for correlation with medicinal species. The PGLS was conducted in R using the package caper (Orme et al., 2012).

Analysis was constrained to one region to mitigate any potential bias due to some regions having a more complete ethnobotanical record in literature than others. North America is an ideal choice because, for one, it has a relatively comprehensive ethnobotanical record, thanks in part to NAEB. However, the general population is, for the most part, not reliant on traditional plant medicine and there is less emphasis on native medicinal plant research here than in other regions with a more complete ethnobotanical record, such as southern and southeastern Asia (Moerman, 2003). This means that species are less likely to be collected or observed specifically because they are medicinal, and the number of occurrence records can be assumed to be an accurate representation of apparency.

## RESULTS

### Global analysis

A total of 1781 categorized medicinal uses matching 465 species across 11 Caryophyllales families were retrieved in the literature search. Of those, 319 species with 1365 uses overlapped with the guide tree (69% species overlap). The species not on the guide tree were discarded from further analysis.

The most uses occurred in the Digestive System Disorders category (Appendix S1; see Supplemental Data with this article). North America had the most species with reported uses (108 species), however, the most reported uses across species and category comes from Southern Asia (392 uses; Appendix S2). Eastern Asia had the highest average number of different use categories per species (4.28; Appendix S2).

Clades with hot nodes were largely consistent between categories, although categories with fewer reported uses predictably had fewer hot nodes. Cactaceae, a betalain-producing family with 1108 species represented on the guide tree, consistently had scattered cold nodes and only a few, mostly deep, hot nodes across the categories, with the notable exception of the Endocrine System Disorder category, which had a hot clade in the *Opuntia* genus of Cactaceae (Appendix S3; Appendix S4). Outside of Cactaceae, cold nodes were relatively uncommon. On the other end of the spectrum, anthocyanin-producing Polygonaceae, with only 565 species represented in the guide tree, was fairly consistently overrepresented across medicinal use categories (Appendix S3, Fig. 2). Hot nodes in Polygonaceae mostly occurred in the clades representing the Rumiceae and Persicarieae. Hot nodes in Caryophyllaceae were few and less consistent across categories, with medicinal uses scattered across the family (Fig. 2).

**Fig. 2.**
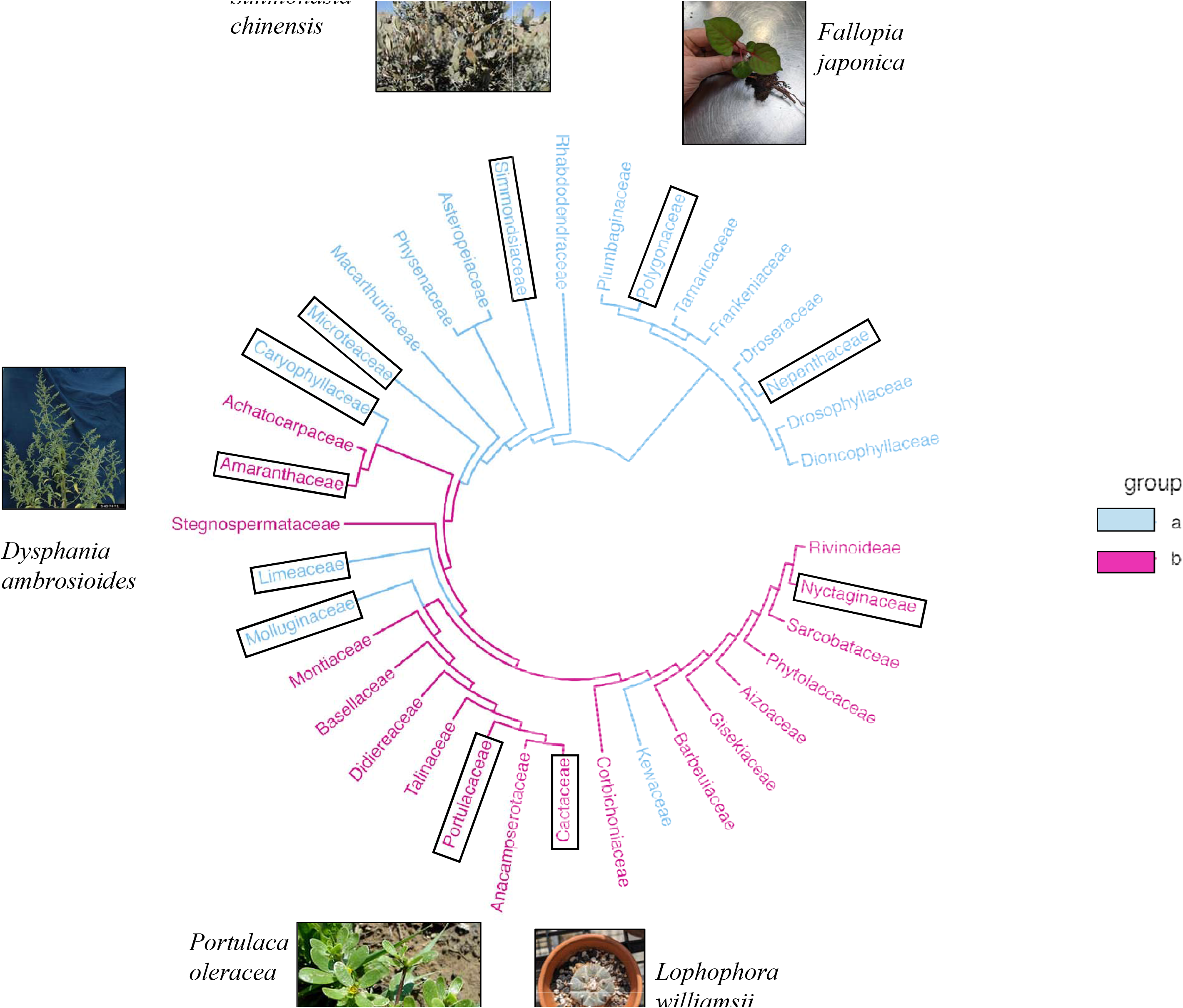
Medicinal use in four major medicinal categories in representative families of Caryophyllales across the globe. Red nodes indicate clades where medicinal use is phylogenetically clustered in that category. Gray nodes indicate clades where medicinal use is underrepresented. The geographical region(s) where a medicinal species is used in that category is given in concentric rings outside the phylogeny. The outermost ring of magenta and blue indicate clades that are either anthocyanin or betalain producing. Although only the top three most-reported categories and the Mental/Nervous System category are shown here, the other, less abundant medicinal categories show the same pattern of hot nodes in Polygonaceae and Amaranthaceae and cold nodes in Cactaceae (with the exception of Endocrine System Disorders; Appendix S4).

Amaranthaceae hot nodes mostly occurred in the Amaranthoideae and Gomphrenoideae subfamilies, with less consistent hot nodes in the Chenopodioideae subfamily occurring in some categories (Fig. 2; Appendix S4).

Due to the low number of reported uses in the Nervous System and Mental Disorders categories and their action on the same body system, the two categories were combined for analysis with Phylocom. All other categories were kept separate.

### North American Analysis

Based on GBIF data, the guide tree was trimmed to 914 species, 103 of which had medicinal use in at least one category, for a total of 300 medicinal uses. Similar patterns of use were observed, although hot clades were less consistently present across categories due to the reduced sample size. When present in Amaranthaceae and Polygonaceae, they were in the same clades highlighted in the global analysis. Nyctaginaceae, now a larger percentage of the tree, also had hot nodes in several categories on the North American tree. Cold nodes overall were less common in the North American analysis, although Cactaceae still had few, inconsistent hot nodes across categories, again with the exception of the “Endocrine System Disorders” category (Fig. 3; Appendix S5). Medicinal use was significantly correlated with a larger apparency (F = 171.2, df = 1, P<2.2e-16, λ = 0.005; Fig. 4).

**Fig. 3.**
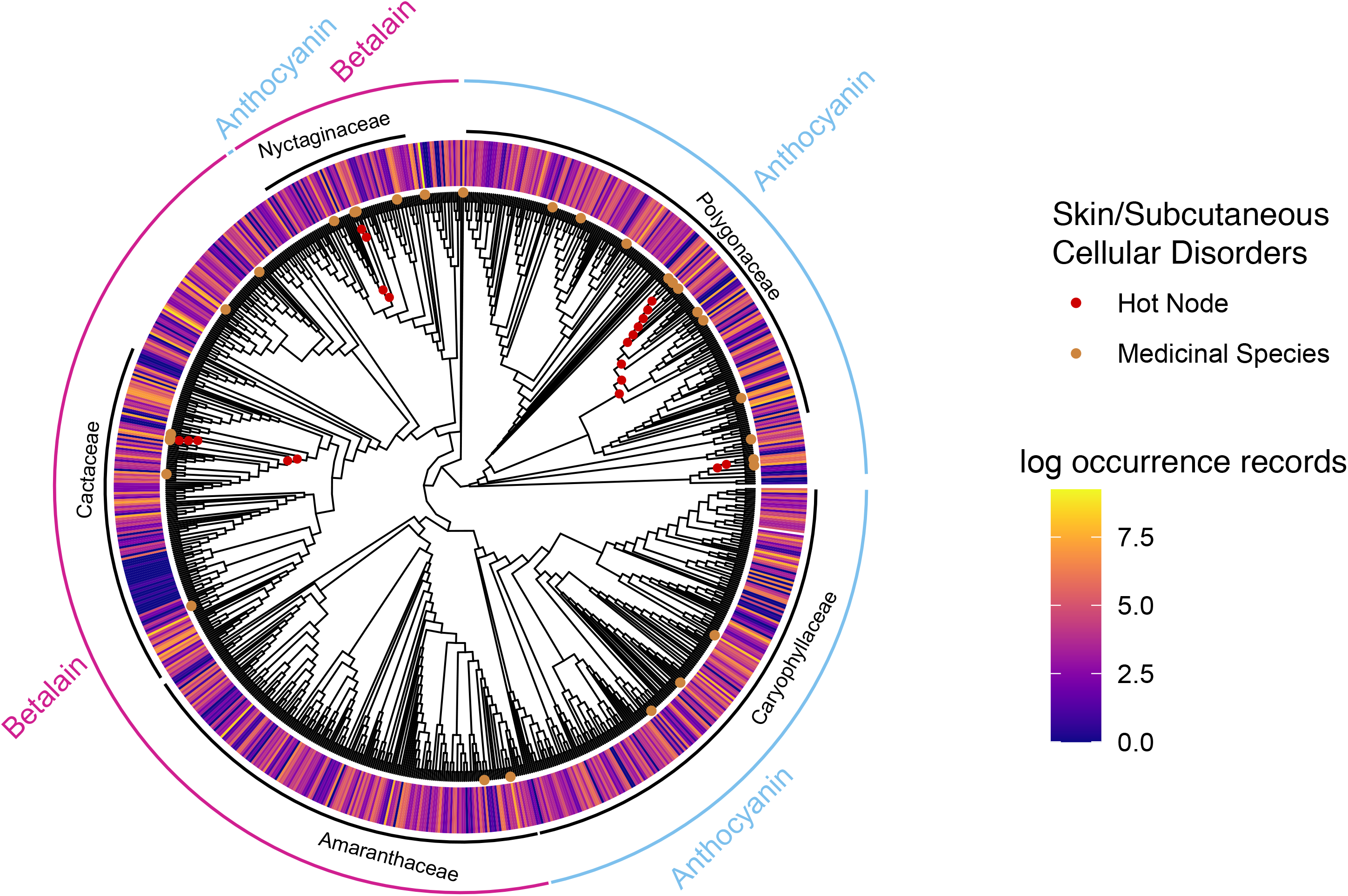
A hot nodes analysis of Skin/Subcutaneous Cellular Tissue Disorders medicinal use in the Caryophyllales species of North America. The ring outside the phylogeny represents log-transformed occurrences from GBIF.

**Fig. 4.**
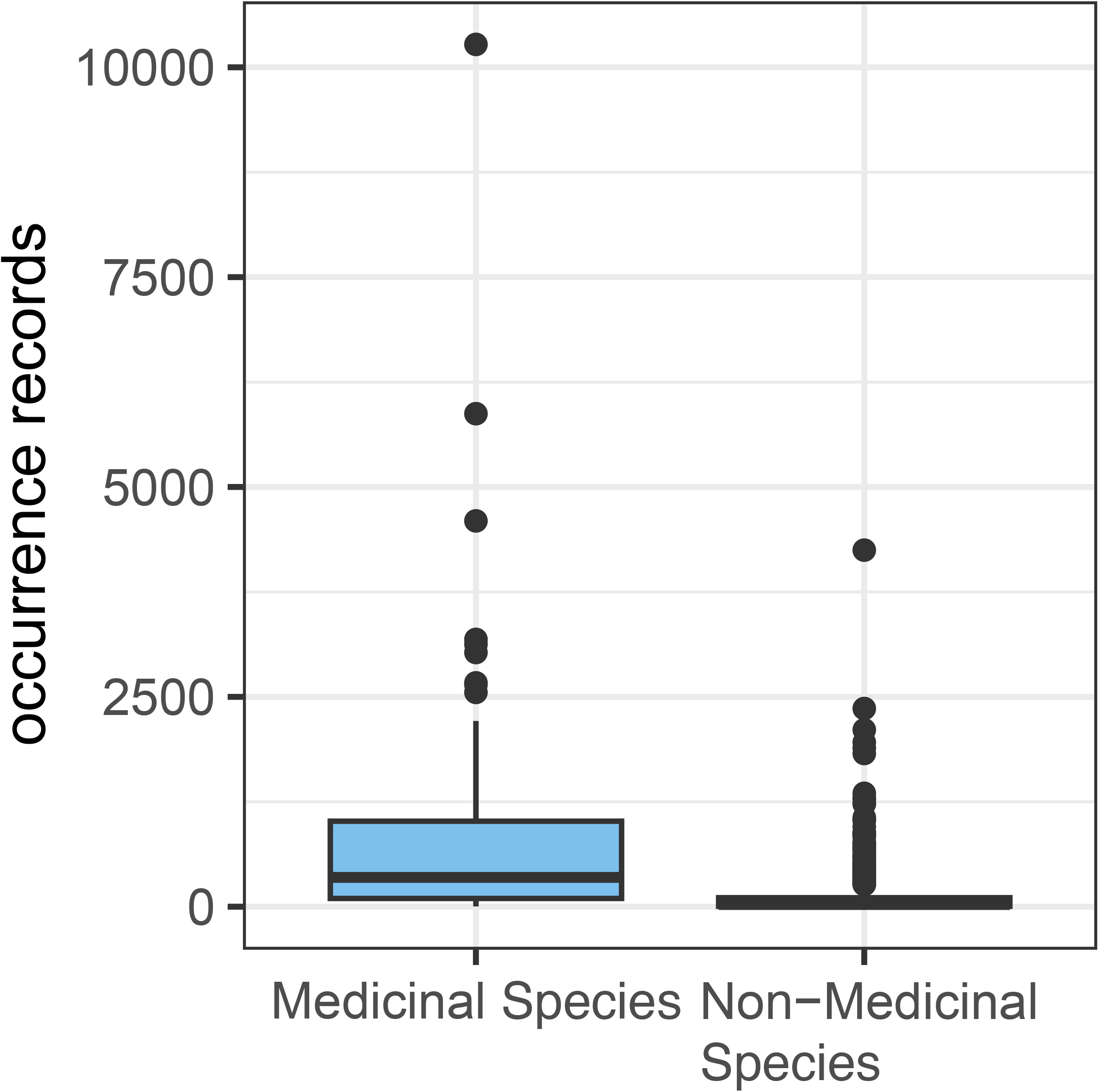
Apparency (as represented by occurrence data from GBIF) of species with any type of medicinal use vs. those with no reported medicinal use.

## DISCUSSION

Despite the dichotomy of Tyr or Phe-dominant metabolism in Caryophyllales, overall, reported traditional medicinal uses did not differ by metabolism type. Instead, the same clades were largely selected for medicinal use repeatedly across all categories and geographic regions. In the global analysis, the two main hot clades in Polygonaceae and Amaranthaceae were also notable for having a pattern of use across many regions in all categories.

Species in these hot clades also have a tendency to be common or even weedy and have a wider geographic range of distribution. Common species being disproportionately used medicinally has been observed in several cultures and locations outside of this work as well (Lewu and Afolayan, 2009; Stepp and Moerman, 2001). For example, the Polygonaceae genus *Rumex*, which is consistently overrepresented across medicinal use categories, is worldwide in distribution, with several species considered invasive in the U.S. In the larger categories, such as Digestive System Disorders, most *Rumex* species are also used in multiple regions. *Rumex* species used in multiple categories also are usually used in multiple regions depending on category (i.e. the ubiquitous *Rumex crispus*). On the other hand, Cactaceae, which had few hot nodes, has many slow-growing species largely restricted to North and South America. Although a few *Opuntia* species have been distributed worldwide by humans and are used outside their native range, for the most part, Cactaceae medicinal use is restricted to the North America and Latin America/Caribbean regions.

### The North American analysis supports apparency as a driver of medicinal plant selection

The pattern of higher medicinal use in these weedier, widespread clades suggests that plant species that have more contact with humans and a variety of human cultures are more likely to be selected for medicinal use across categories than the species less apparent to humans. This is supported by the additional analysis restricted to North America. While hot nodes are less consistent across medicinal use categories in North America, which isn’t surprising given the reduced sample size, the same clades in Polygonaceae and Amaranthaceae tend to either have hot nodes or be one of the few clades with medicinal use of any kind. The strong case for apparency being the leading factor driving human medicinal plant selection is made by the preference of certain clades across North American cultures, attribution of many medicinal uses across categories to the same clades, and the significant correlation between occurrence data and whether or not a plant is used medicinally.

### The availability and resource availability hypotheses

Apparency as a main driver of selection is consistent with the availability hypothesis and previous studies finding that weedy species are more likely to be selected for medicinal use due to frequent human contact (Gaoue et al., 2017; Stepp and Moerman, 2001; Voeks, 2004). In fact, weedy species tend to follow humans as they move across the globe, increasing the likelihood of contact with different cultural groups, and helping them acquire uses in new categories (Chapman et al., 2017). This hypothesis also explains the lack of use in Cactaceae, which is restricted to North and South America, with the exception of several widely introduced species (i.e. *Opuntia* spp.). Tellingly, the species of Cactaceae used in the most categories (15) is *Opuntia ficus-indica,* which also sees use in the most regions (6). Additionally, as Stepp and Moerman (2001) point out, humans are more likely to seek out medicines that are abundant and easily acquired when necessary, rather than relying on uncommon or difficult-to-access plants. While the availability hypothesis argues that underlying metabolism is not the primary factor underlying human selection for medicinal use, there may be other factors to consider.

There is prior evidence to suggest that weedy species, because they are faster growing and shorter lived, tend to have more quickly-made, bioactive, and therefore potentially medicinal, compounds rather than investing in longer-lasting, but slower and costly to produce defenses such as lignins and tannins (Endara and Coley, 2011). This is in line with the resource availability hypothesis, which posits that plants adapted to high-resource areas, such as disturbed areas, tend to be quick-growing and therefore put more resources into “quick” herbivory defenses, like bioactive specialized metabolites, rather than “slow” defenses that reduce the plant’s digestibility (Endara and Coley, 2011; Gaoue et al., 2017). The weedy species, then, may have been selected by humans because they truly do have more medicinal use than their non-weedy relatives. This would also explain the relative lack of medicinal uses in Cactaceae, as a comparatively slow-growing group that may have more investment in slow defenses, such as spines and glochids, than bioactive SMs. The patterns of medicinal use most likely are caused by a combination of both apparency and SM, where weedy species have evolved to have more SMs to deter herbivory, and also are more available to be selected for medicinal use.

While statistical power is limited in medicinal categories with few uses, and it is difficult to separate out the two potential drivers of selection, there may still be some signal of metabolite-based selection under the overwhelming signal of apparency-driven selection. It is compelling to note that the global Mental/Nervous System Disorders category has few uses in Polygonaceae (an anthocyanin family), and the only two hot nodes in the family occur at the tips (Figure 2). Although a few of the other small categories (i.e. Sensory System Disorders and Pregnancy/Birth/Puerperium Disorders) also lose this hot clade, they typically show little clustering in Amaranthaceae (Appendix S4). In the Mental/Nervous System Disorders Category, the Amaranthaceae hot clade (a betalain lineage) still shows clustering. Further, in the North American analysis of Mental and Nervous System Disorders, there is not a single medicinal use in anthocyanin lineages, with most occurring in either Nyctaginaceae, Amaranthaceae, or scattered through Cactaceae. Despite the limited uses reported for these categories, the even more limited use in anthocyanin lineages is suggestive of the predicted pattern of more use in lineages with Tyr-dominant metabolism. A more targeted exploration of use of Caryophyllales species in this category is needed to definitively distinguish this pattern from statistical noise.

### Caveats

The ethnobotanical record available in literature is likely incomplete and, based on the geographic patterns observed in Caryophyllales, somewhat biased. It is telling that the most reported uses come from Eastern Asia, where Traditional Chinese Medicine has been actively studied and documented for some time, and the most medicinal species come from North America, which has an extensive database of traditional use (Moerman, 2003). If more ethnobotanical research were done in less studied areas, such as Oceania, which has native and introduced species from several of the families reviewed here, but only four medicinal species recovered in our literature search, it is possible that we would see differences start to emerge between medicinal categories. However, the fact that the hot clades are clades with wide geographic distributions suggest that, despite more documentation of medicinal use in some regions, these clades would still be consistently hot across categories. Additionally, the North American analysis, while having less consistently present hot nodes, did not show a radically different pattern of where the hot nodes were placed when present.

In the guide tree used, deep relationships in Cactaceae were poorly supported.

Phylocom’s nodesig command is unable to account for either branch length or phylogenetic uncertainty, so hot node estimation may be biased by a poorly supported tree, However, given the overall limited medicinal use reported in Cactaceae and that 38.7% of the family’s medicinal species occur in one genus, *Opuntia*, the lack of hot nodes in the family is unlikely to be a result of phylogenetic uncertainty in the family.

The selection by apparency rather than by dominant metabolism type may be driven by the availability or resource availability hypotheses as explained above, or, alternatively, there may be overlap between the bioactivity of the metabolites derived from the two related amino acids. Phenylalanine-derived specialized metabolism includes phenolics like flavonoids, coumarins, and lignans, which have been ascribed a wide range of potential health benefits.

While tyrosine-derived metabolites tend to be less ubiquitous, some of the best-known include neurologically-active compounds like mescaline, dopamine, and morphine (Xu et al., 2019).

However, other Tyr-derived compounds may contribute to Tyr metabolism having as broad action as Phe metabolism. For example, apart from neurological effect, some Tyr-derived catecholamines like epinephrine act on the cardiovascular and respiratory system (Abul-Ainine, 2002; Bao et al., 2007). If Tyr- and Phe-derived metabolism overall have enough overlap in medicinal activity so as to be indistinguishable from each other, human selection of apparent plants would still be driven by the resource availability hypothesis, and dominant metabolism type would be entirely irrelevant in driving human selection.

## CONCLUSIONS

Given the renewed interest in natural products for drug discovery, understanding the relationships between specialized metabolism, phylogeny, and medicinal use is important in prioritizing plants to investigate and narrowing down the metabolites responsible for medicinal function. In addition, an approach combining phylogenetic, metabolic, and global use information can elucidate cross-cultural patterns of medicinal use and what is driving them.

Here, we show that, at least in Caryophyllales, the apparency of a plant to humans is a dominant factor in its being selected for medicinal use across cultures, as opposed to the plant’s broad SM profile. Although there are several possible explanations for this, a combination of humans selecting readily apparent plants (availability hypothesis) and those same plants having greater bioactive activity (resource availability hypothesis) appear to be the most likely. Further investigation of the Tyr- and Phe- derived metabolic diversity in both medicinal and non-medicinal clades of the order would help determine which or both of these hypotheses is driving selection of medicinal plants in Caryophyllales and help untangle the effects of apparency from other potential drivers of medicinal plant selection.

## ACKNOWLEDGMENTS

We thank M. Mohamed and L. Magpali for their assistance in data collection, and A. Lee, R.A. Mohn, and A. Smeenk for their feedback on the work. Funding was provided by the National Science Foundation (Award#1939226).

## AUTHOR CONTRIBUTIONS

A.C. conceived of the research, collected and analyzed data, and wrote the manuscript, L.P. contributed to the design of the research and manuscript revision. L.B. wrote scripts for data collection. Y.Y. aided in conception and design, and manuscript revision.

## SUPPORTING INFORMATION

Additional supporting information may be found online in the Supporting Information section at the end of the article.

Appendix S1 - Table with medicinal use counts by category Appendix S2 - Table with medicinal use by region

Appendix S3 - Table with medicinal use by family

Appendix S4 - Hot and cold nodes mapped out on a phylogeny of selected Caryophyllales families across the globe. All medicinal categories not part of Fig. 2 are included.

Appendix S5 - Hot and cold nodes mapped out on a phylogeny of North American Caryophyllales species. A selection of medicinal categories is included.

