## Supplemental 1-3 for "Traditional medicinal use is linked with apparency, not specialized metabolite profiles in the order Caryophyllales"

| **Category of Use** | Abnormalities | Blood System Disorders | Circulatory System Disorders | Digestive System Disorders | Endocrine System Disorders | Genitourinary System Disorders | Ill-Defined Symptoms | Immunological Disorders | Infections and Infestations | Inflammation |
|---|---|---|---|---|---|---|---|---|---|---|
| **Total # of Species with Reported Use** | 27 | 29 | 66 | 156 | 39 | 114 | 7 | 3 | 165 | 72 |
| **% of Medicinal Species with Use in this category** | 8.46% | 9.09% | 20.69% | 48.9% | 12.23% | 35.74% | 2.19% | 0.09% | 51.72% | 22.57% |

| **Category of Use** | Injuries | Mental Disorders | Metabolic System Disorders | Muscular-Skeletal System Disorders | Neoplasms | Nervous System Disorders | Nutrition Disorders | Pain | Poisonings | Pregnancy, Birth, and Puerperium Disorders | Respiratory System Disorders |
|---|---|---|---|---|---|---|---|---|---|---|---|
| **Total # of Species with Reported Use** | 93 | 32 | 21 | 68 | 24 | 20 | 59 | 72 | 56 | 25 | 77 |
| **% of Medicinal Species with Use in this category** | 29.15% | 10.03% | 6.58% | 21.31% | 7.52% | 6.27% | 18.50% | 22.57 % | 17.55% | 7.83% | 24.13% |

| **Category of Use** | Sensory System Disorders | Skin and Subcutaneous Cellular Tissue Disorders |
|---|---|---|
| **Total # of Species with Reported Use** | 28 | 112 |
| **% of Medicinal Species with Use in this category** | 8.78% | 35.11% |

**Appendix S1-**Number of reported uses from each medicinal category

**Appendix S2**

| **Region** | Sub-Saharan Africa | Northern Africa | Central Asia | Eastern Asia | Western Asia | Southern Asia | Southeastern Asia | Oceania | Europe | Latin America & the Caribbean | North America |
|---|---|---|---|---|---|---|---|---|---|---|---|
| **Number of Categorized Uses** | 138 | 67 | 11 | 291 | 116 | 392 | 57 | 9 | 96 | 220 | 295 |
| **Number of Medicinal Species** | 41 | 23 | 5 | 68 | 45 | 102 | 20 | 4 | 29 | 60 | 108 |
| **Different uses per plant species** | 3.28 | 2.91 | 2.2 | 4.28 | 2.32 | 3.84 | 2.85 | 2.25 | 3.31 | 3.67 | 2.73 |

**Appendix S3**

| **Family** | Polygonaceae | Simmondsiaceae | Microteaceae | Caryophyllaceae | Amaranthaceae | Limeaceae | Molluginaceae | Cactaceae | Nyctaginaceae | Nepenthaceae | Portulacaceae |
|---|---|---|---|---|---|---|---|---|---|---|---|
| **Number of Categorized Uses** | 433 | 8 | 4 | 243 | 372 | 8 | 27 | 134 | 89 | 21 | 26 |
| **Number of Medicinal Species on guide Tree** | 95 | 1 | 1 | 67 | 72 | 3 | 5 | 44 | 22 | 6 | 3 |
| **Number of Total species on guide Tree** | 565 | 1 | 2 | 787 | 827 | 6 | 40 | 1108 | 144 | 85 | 64 |
