## Supplemental 4 for "Traditional medicinal use is linked with apparency, not specialized metabolite profiles in the order Caryophyllales"

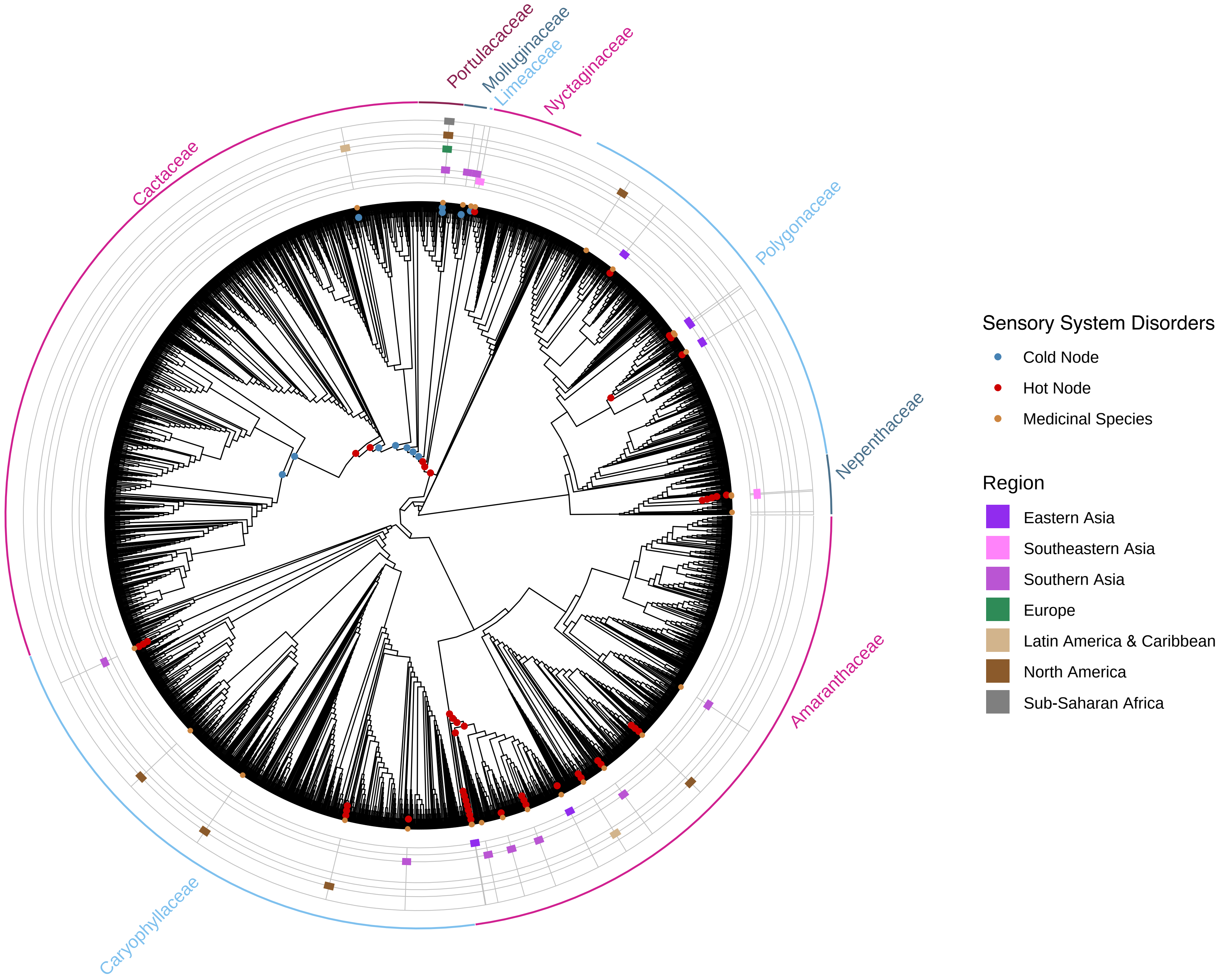

Portulacaceae
Molluginaceae
Limeaceae
Nyctaginaceae
Cactaceae
Polygonaceae
Nepenthaceae
Amaranthaceae
Caryophyllaceae
Sensory System Disorders
Cold Node
Hot Node
Medicinal Species
Region
Eastern Asia
Southeastern Asia
Southern Asia
Europe
Latin America & Caribbean
North America
Sub-Saharan Africa

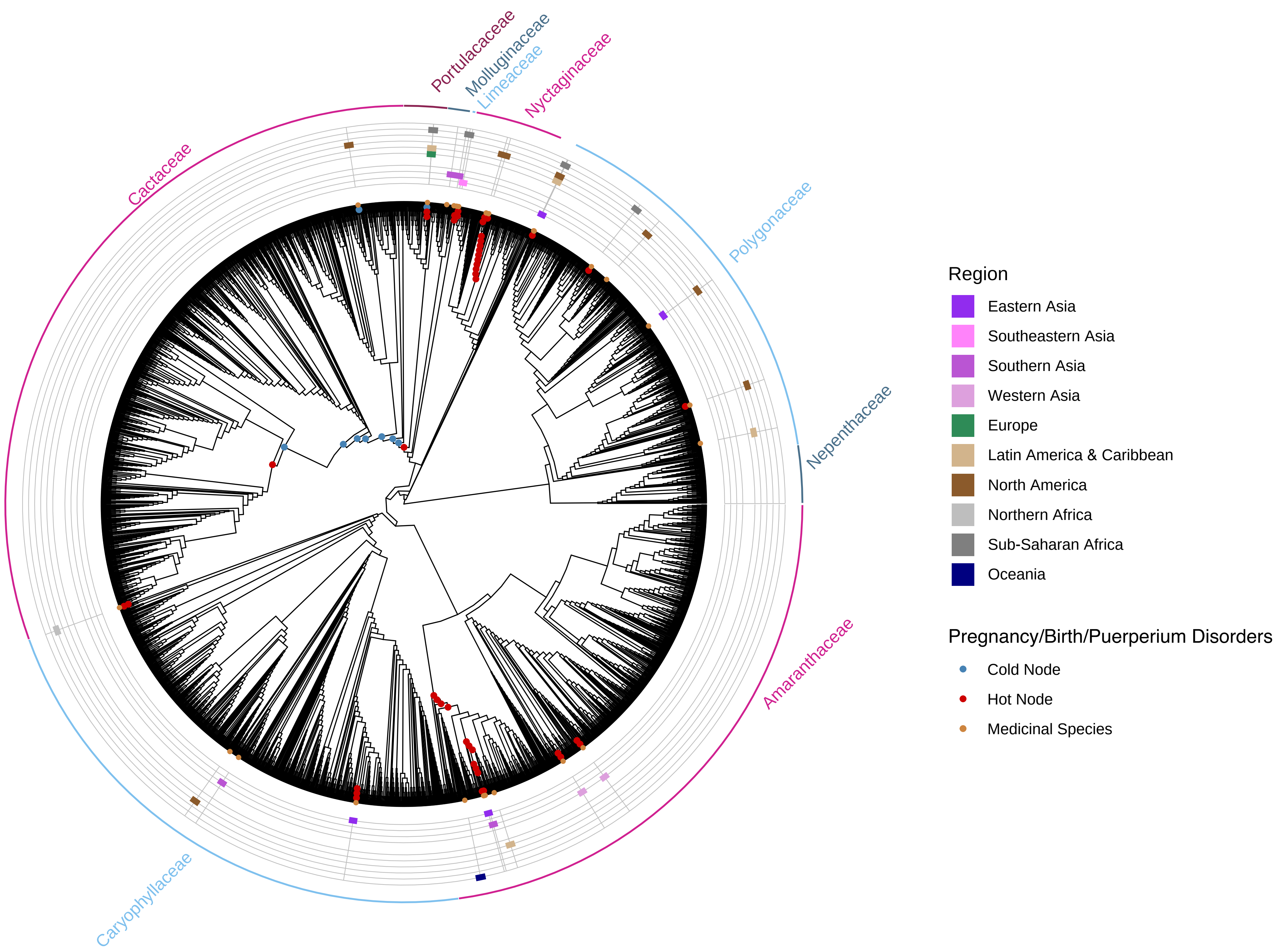

Portulacaceae
Molluginaceae
Limeaceae
Nyctaginaceae
Cactaceae
Polygonaceae
Nepenthaceae
Amaranthaceae
Caryophyllaceae
Region
Eastern Asia
Southeastern Asia
Southern Asia
Western Asia
Europe
Latin America & Caribbean
North America
Northern Africa
Sub-Saharan Africa
Oceania
Pregnancy/Birth/Puerperium Disorders
Cold Node
Hot Node
Medicinal Species

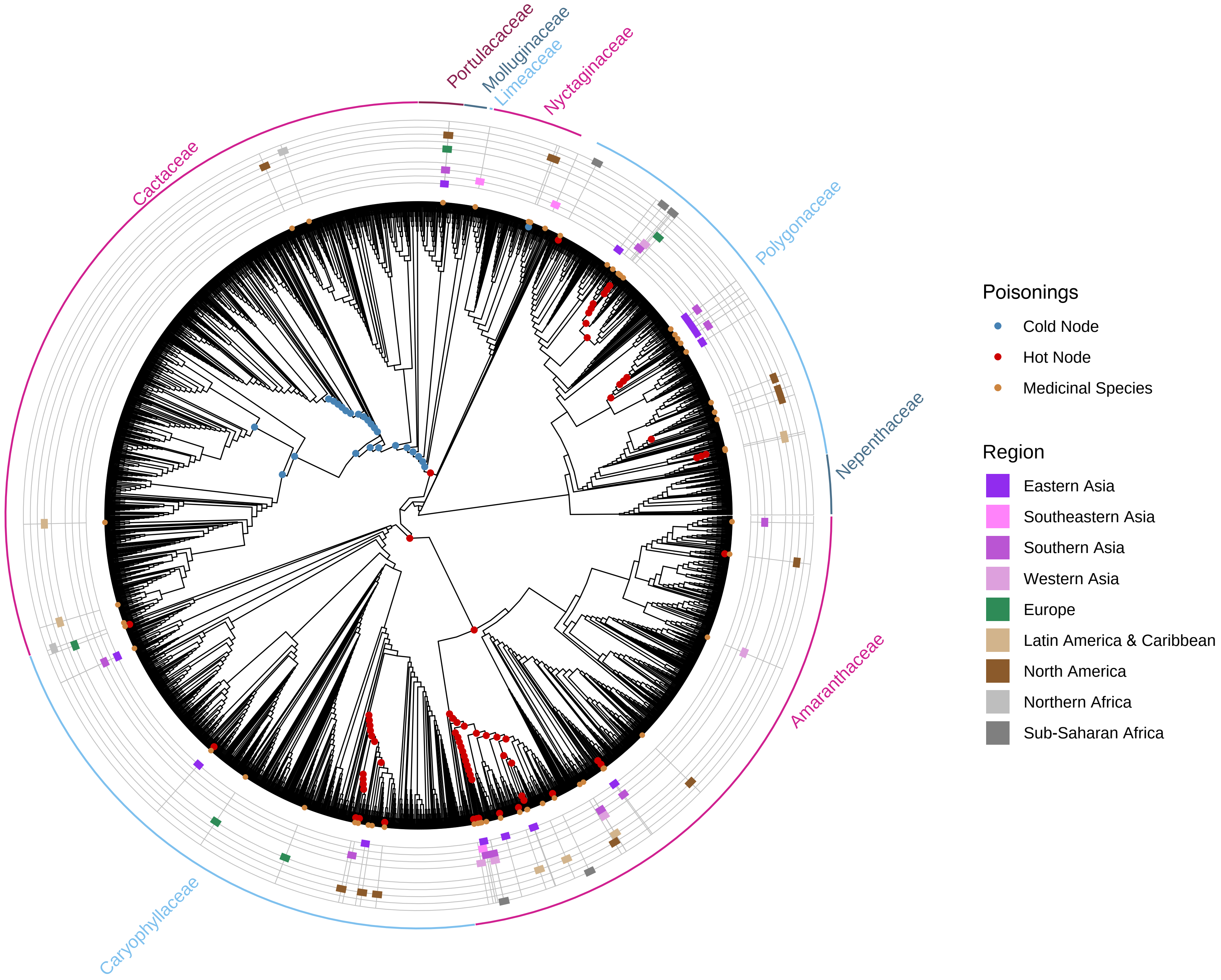

Portulacaceae
Molluginaceae
Limeaceae
Nyctaginaceae
Cactaceae
Polygonaceae
Nepenthaceae
Amaranthaceae
Caryophyllaceae
Poisonings
Cold Node
Hot Node
Medicinal Species
Region
Eastern Asia
Southeastern Asia
Southern Asia
Western Asia
Europe
Latin America & Caribbean
North America
Northern Africa
Sub-Saharan Africa

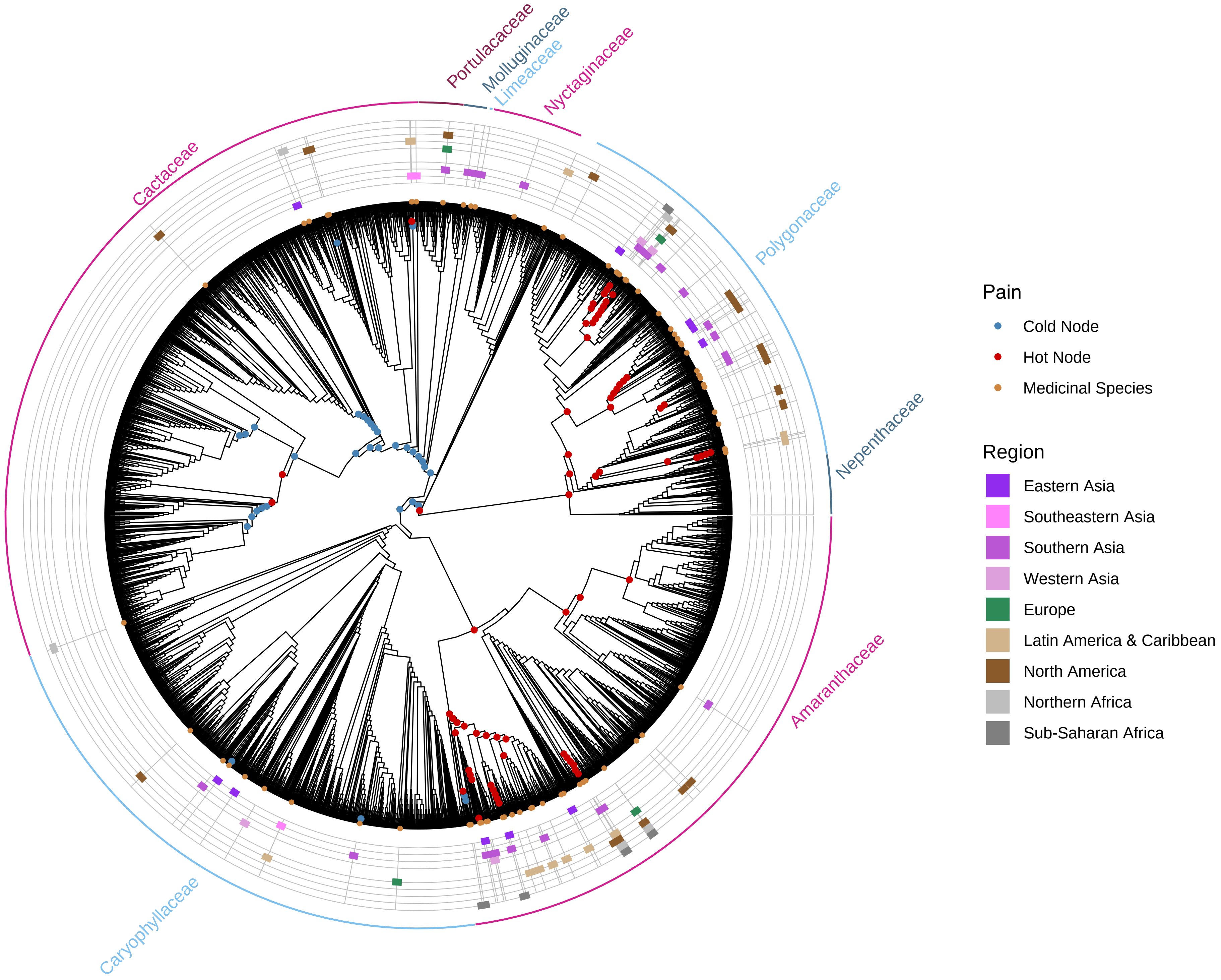

Portulacaceae
Molluginaceae
Limeaceae
Nyctaginaceae
Cactaceae
Polygonaceae
Nepenthaceae
Amaranthaceae
Caryophyllaceae
Pain
Cold Node
Hot Node
Medicinal Species
Region
Eastern Asia
Southeastern Asia
Southern Asia
Western Asia
Europe
Latin America & Caribbean
North America
Northern Africa
Sub-Saharan Africa

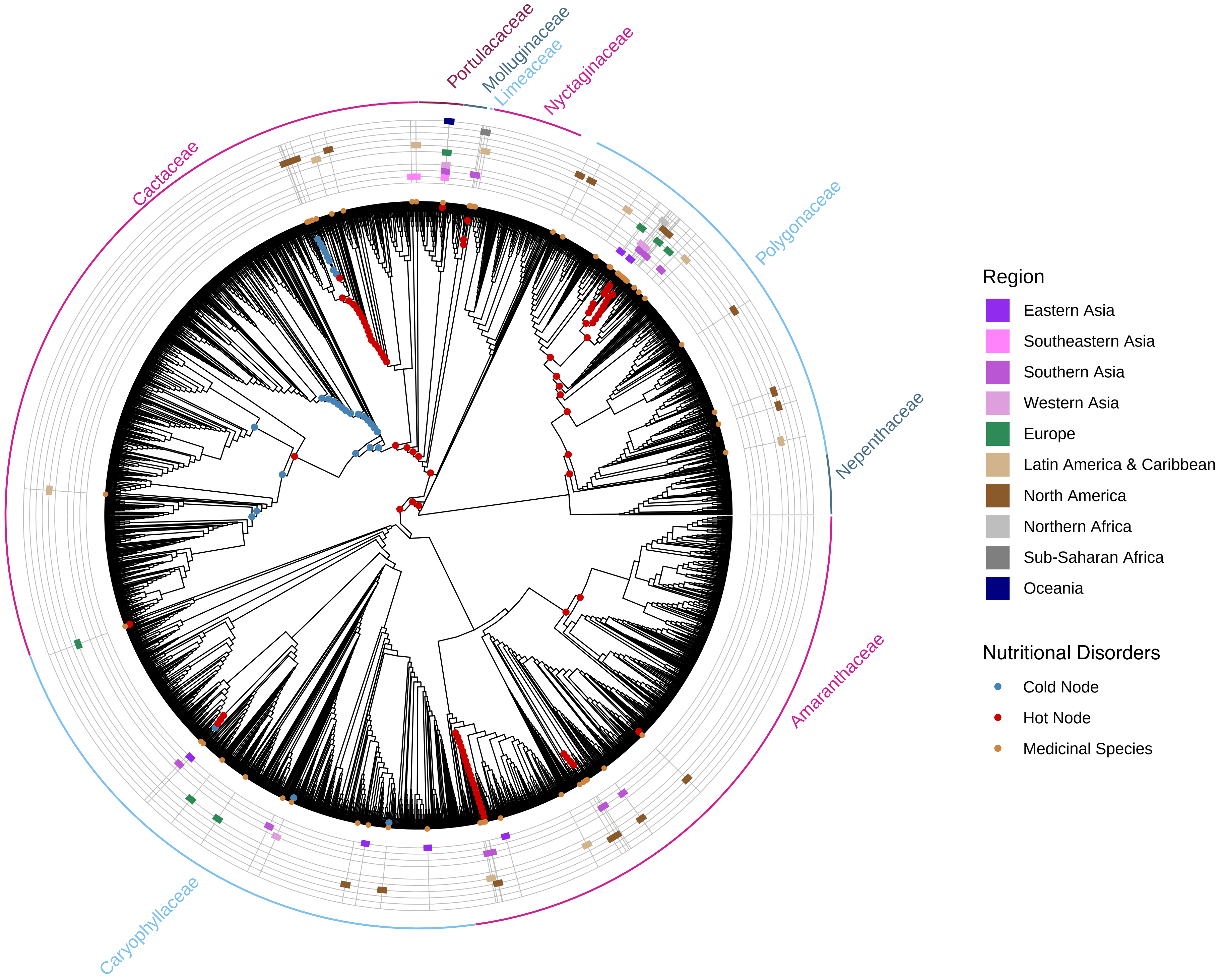
Portulacaceae
Molluginaceae
Limeaceae
Nyctaginaceae
Polygonaceae
Nepenthaceae
Amaranthaceae
Caryophyllaceae
Cactaceae
Region
Eastern Asia
Southeastern Asia
Southern Asia
Western Asia
Europe
Latin America & Caribbean
North America
Northern Africa
Sub-Saharan Africa
Oceania
Nutritional Disorders
Cold Node
Hot Node
Medicinal Species

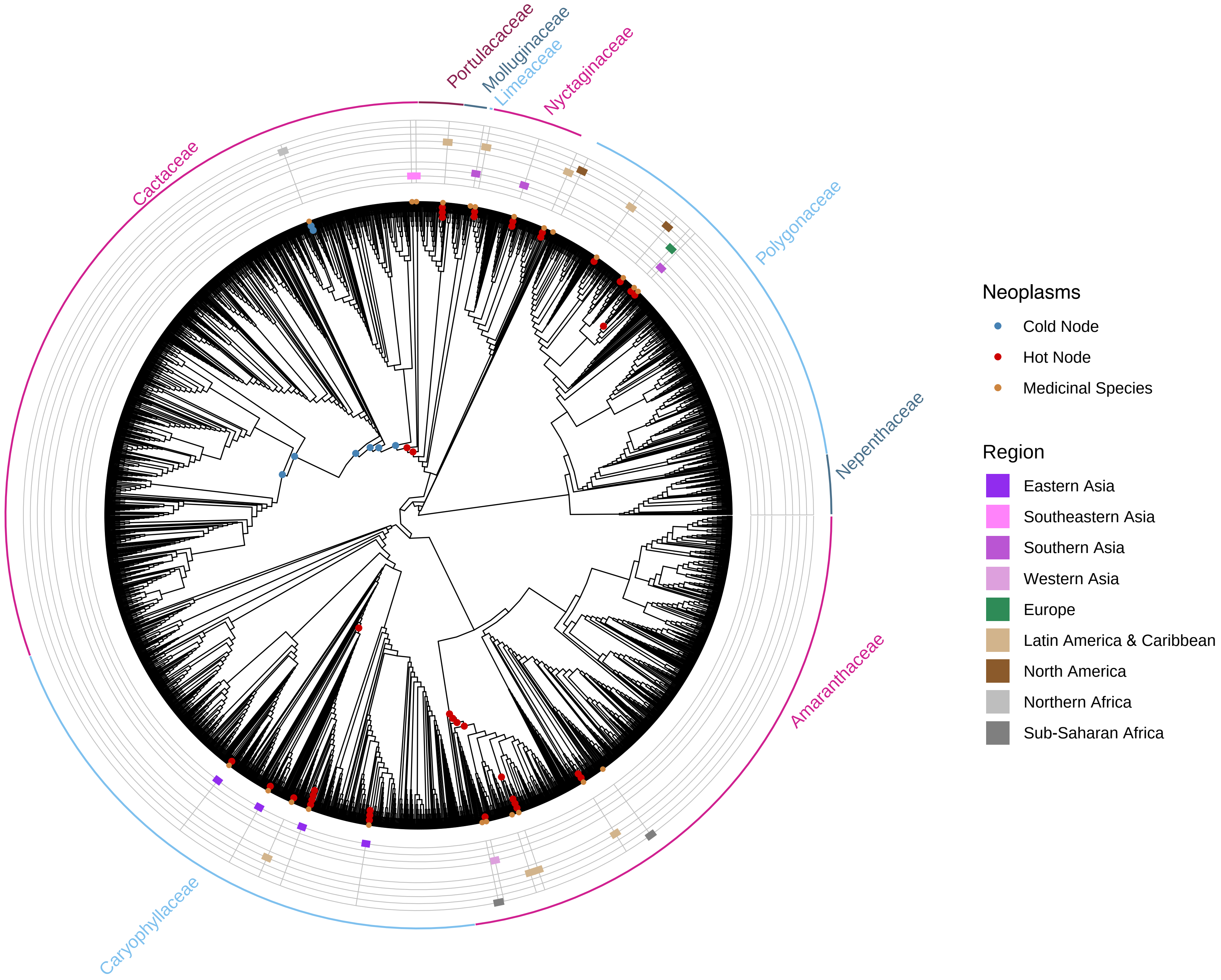
Portulacaceae
Molluginaceae
Limeaceae
Nyctaginaceae
Cactaceae
Polygonaceae
Nepenthaceae
Amaranthaceae
Caryophyllaceae
Neoplasms
Cold Node
Hot Node
Medicinal Species
Region
Eastern Asia
Southeastern Asia
Southern Asia
Western Asia
Europe
Latin America & Caribbean
North America
Northern Africa
Sub-Saharan Africa

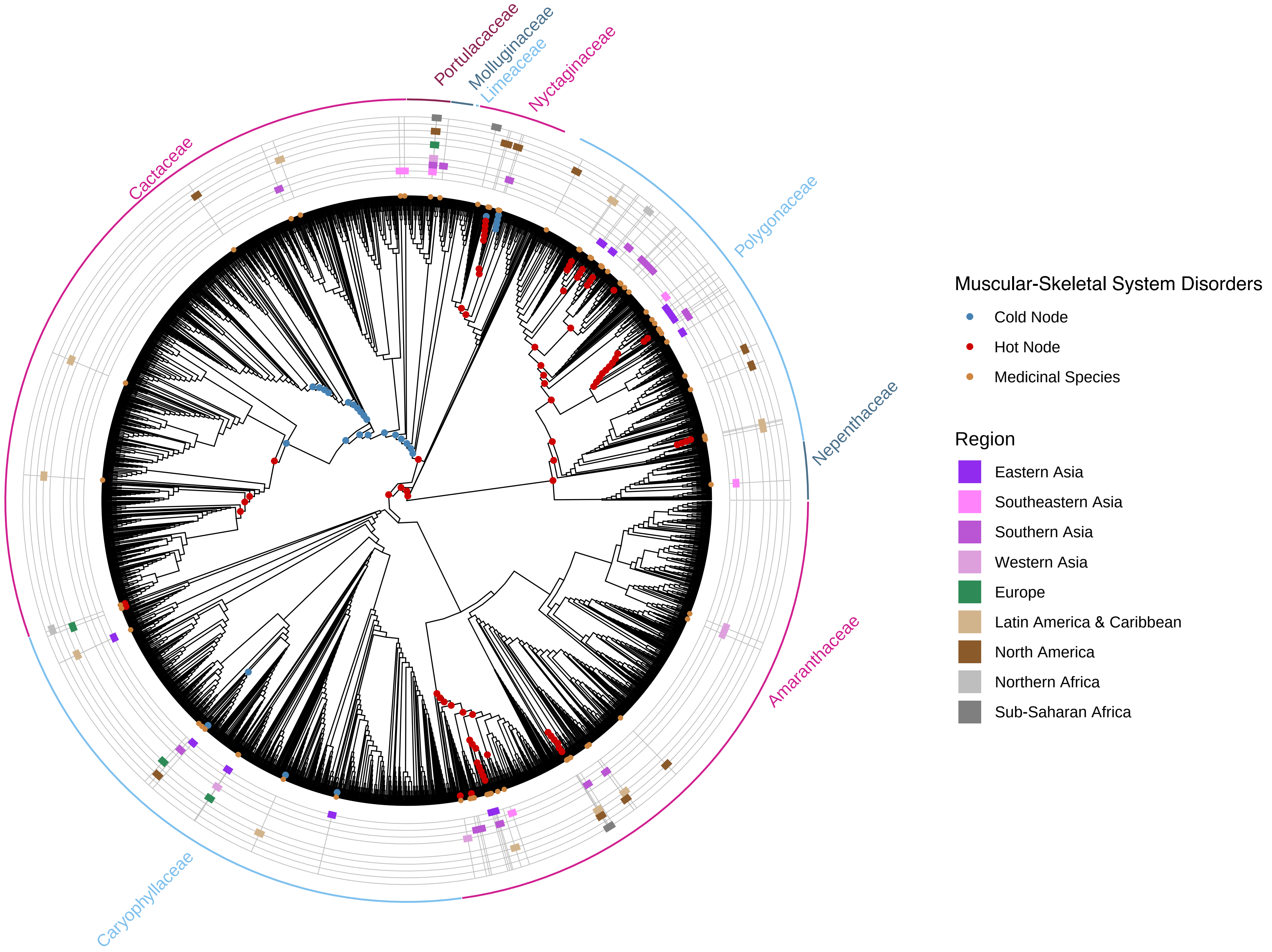
Portulacaceae
Molluginaceae
Limeaceae
Nyctaginaceae
Cactaceae
Polygonaceae
Nepenthaceae
Amaranthaceae
Caryophyllaceae
Muscular-Skeletal System Disorders
Cold Node
Hot Node
Medicinal Species
Region
Eastern Asia
Southeastern Asia
Southern Asia
Western Asia
Europe
Latin America & Caribbean
North America
Northern Africa
Sub-Saharan Africa

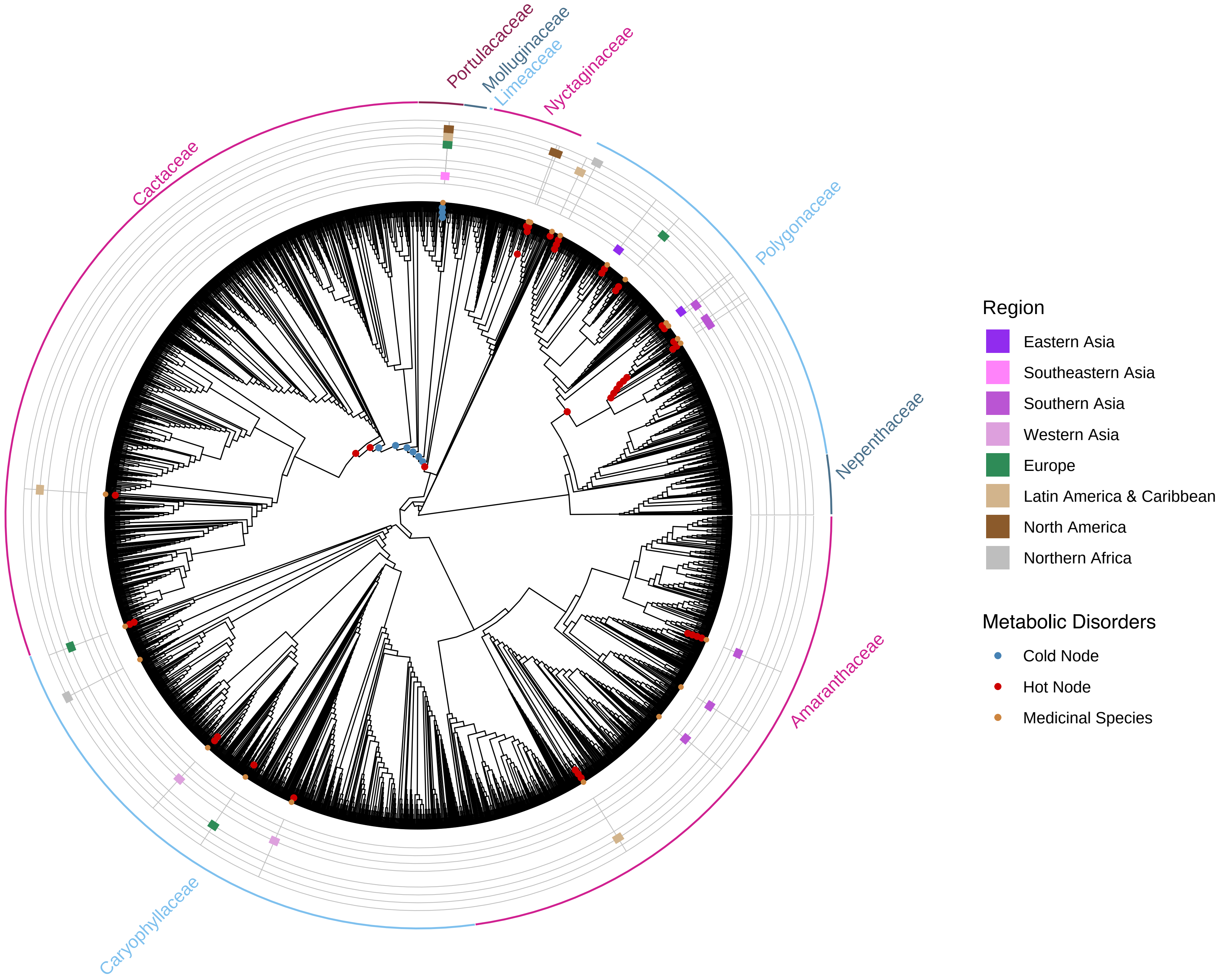

Portulacaceae
Molluginaceae
Limeaceae
Nyctaginaceae
Cactaceae
Polygonaceae
Nepenthaceae
Amaranthaceae
Caryophyllaceae
Region
Eastern Asia
Southeastern Asia
Southern Asia
Western Asia
Europe
Latin America & Caribbean
North America
Northern Africa
Metabolic Disorders
Cold Node
Hot Node
Medicinal Species

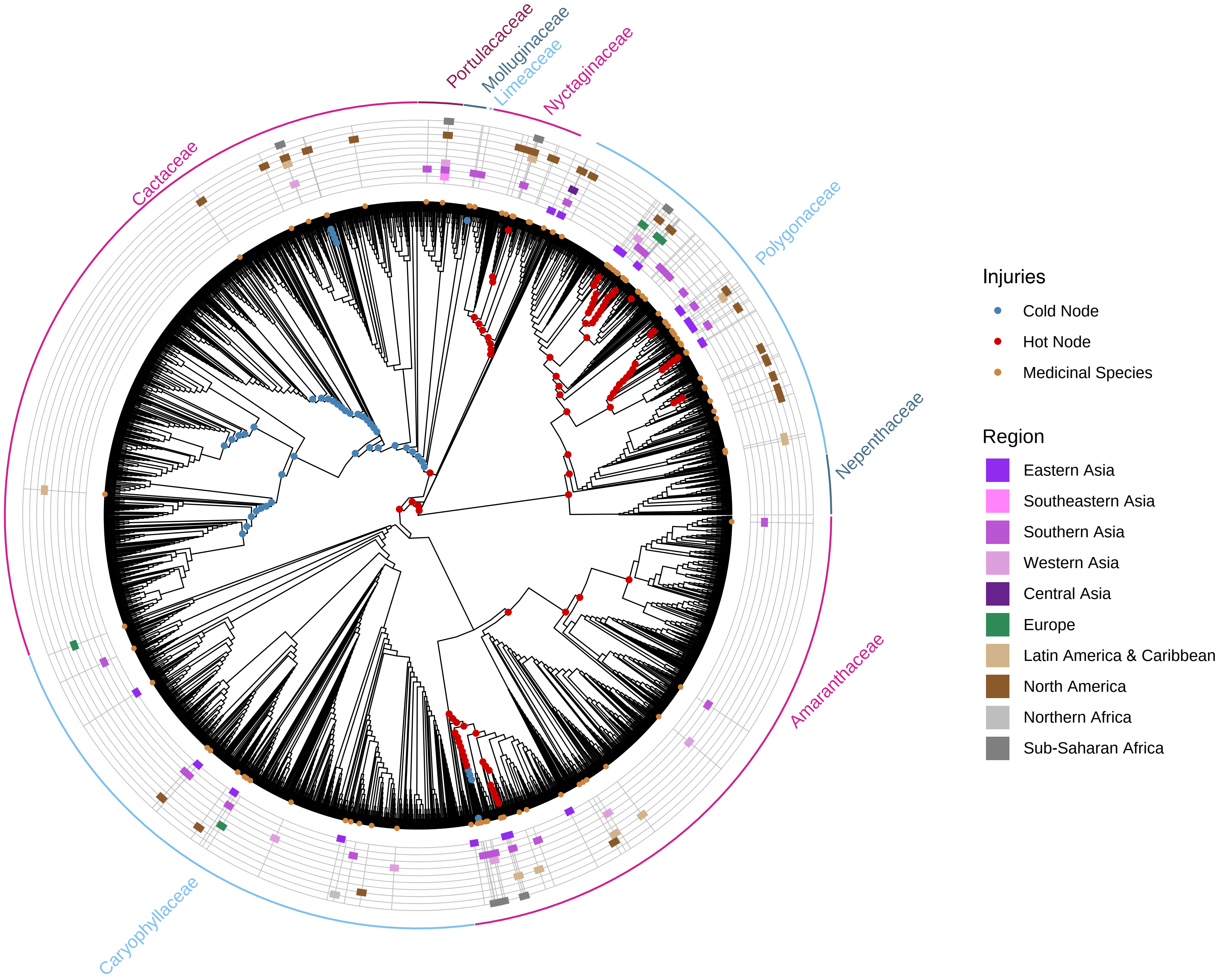

Portulacaceae
Molluginaceae
Limeaceae
Nyctaginaceae
Cactaceae
Polygonaceae
Nepenthaceae
Amaranthaceae
Caryophyllaceae
Injuries
Cold Node
Hot Node
Medicinal Species
Region
Eastern Asia
Southeastern Asia
Southern Asia
Western Asia
Central Asia
Europe
Latin America & Caribbean
North America
Northern Africa
Sub-Saharan Africa

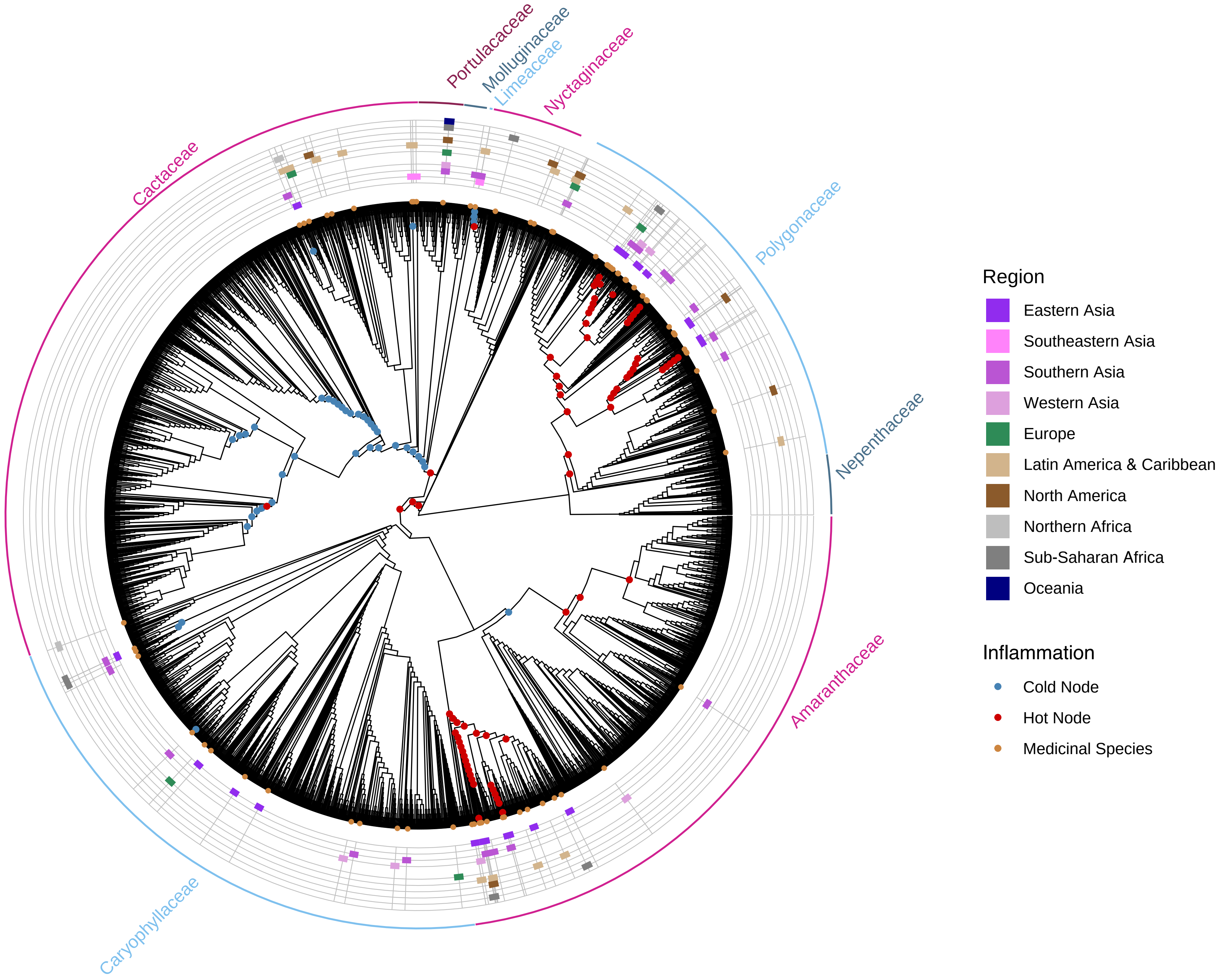
Portulacaceae
Molluginaceae
Limeaceae
Nyctaginaceae
Cactaceae
Polygonaceae
Nepenthaceae
Amaranthaceae
Caryophyllaceae
Region
Eastern Asia
Southeastern Asia
Southern Asia
Western Asia
Europe
Latin America & Caribbean
North America
Northern Africa
Sub-Saharan Africa
Oceania
Inflammation
Cold Node
Hot Node
Medicinal Species

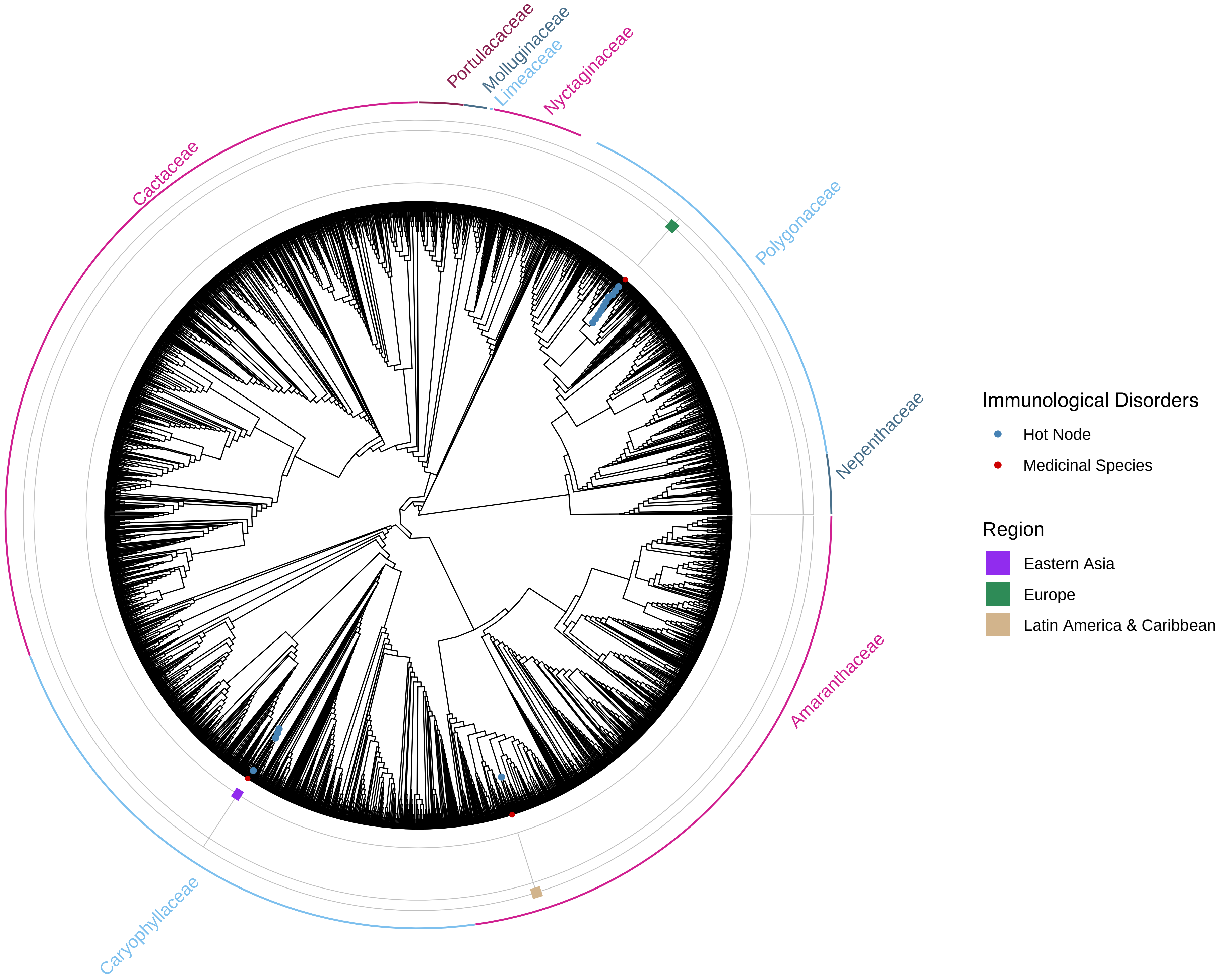

Portulacaceae
Molluginaceae
Limeaceae
Nyctaginaceae
Cactaceae
Polygonaceae
Nepenthaceae
Amaranthaceae
Caryophyllaceae
Immunological Disorders
Hot Node
Medicinal Species
Region
Eastern Asia
Europe
Latin America & Caribbean

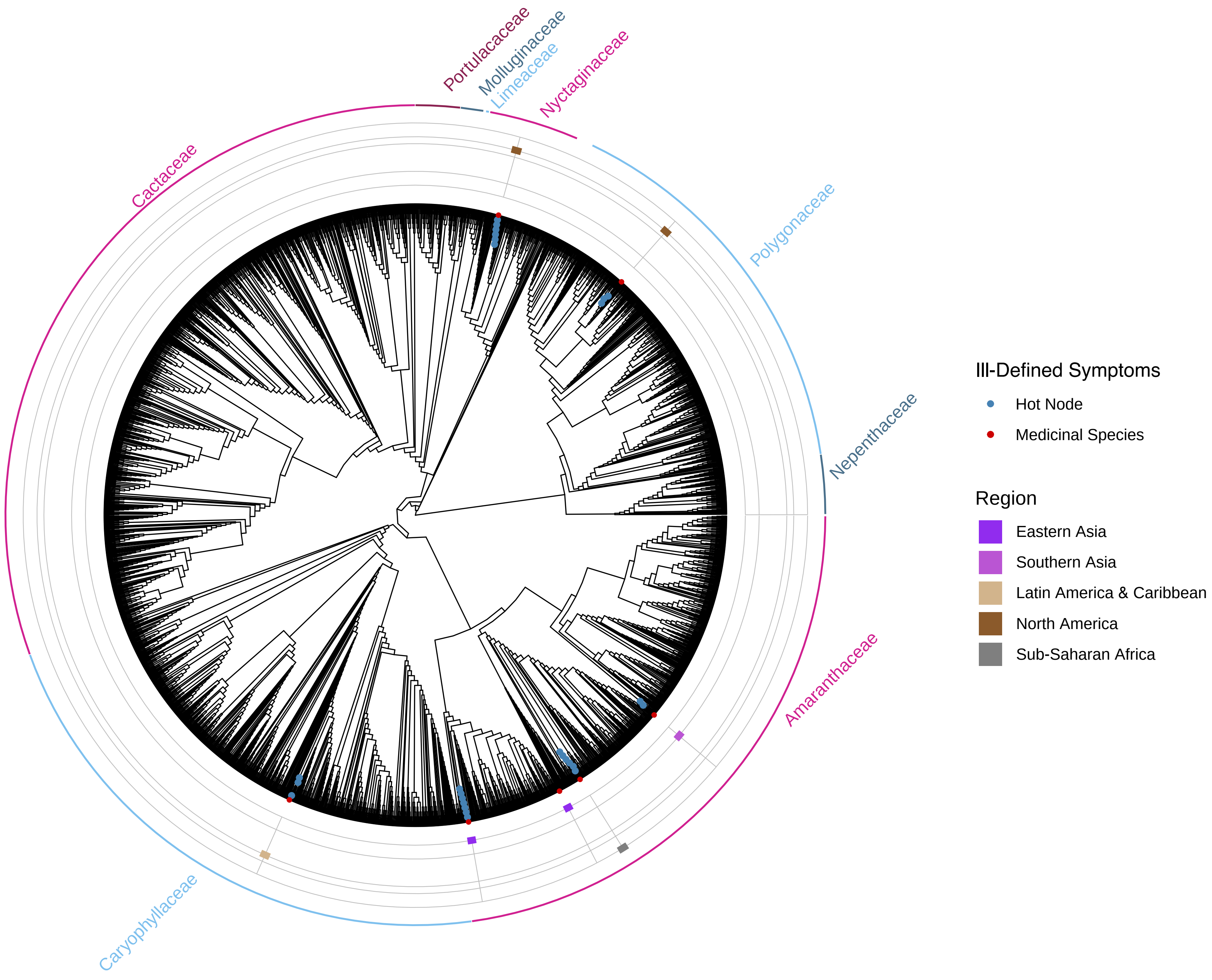

Portulacaceae
Molluginaceae
Limeaceae
Nyctaginaceae
Cactaceae
Polygonaceae
Nepenthaceae
Amaranthaceae
Caryophyllaceae
Ill-Defined Symptoms
Hot Node
Medicinal Species
Region
Eastern Asia
Southern Asia
Latin America & Caribbean
North America
Sub-Saharan Africa

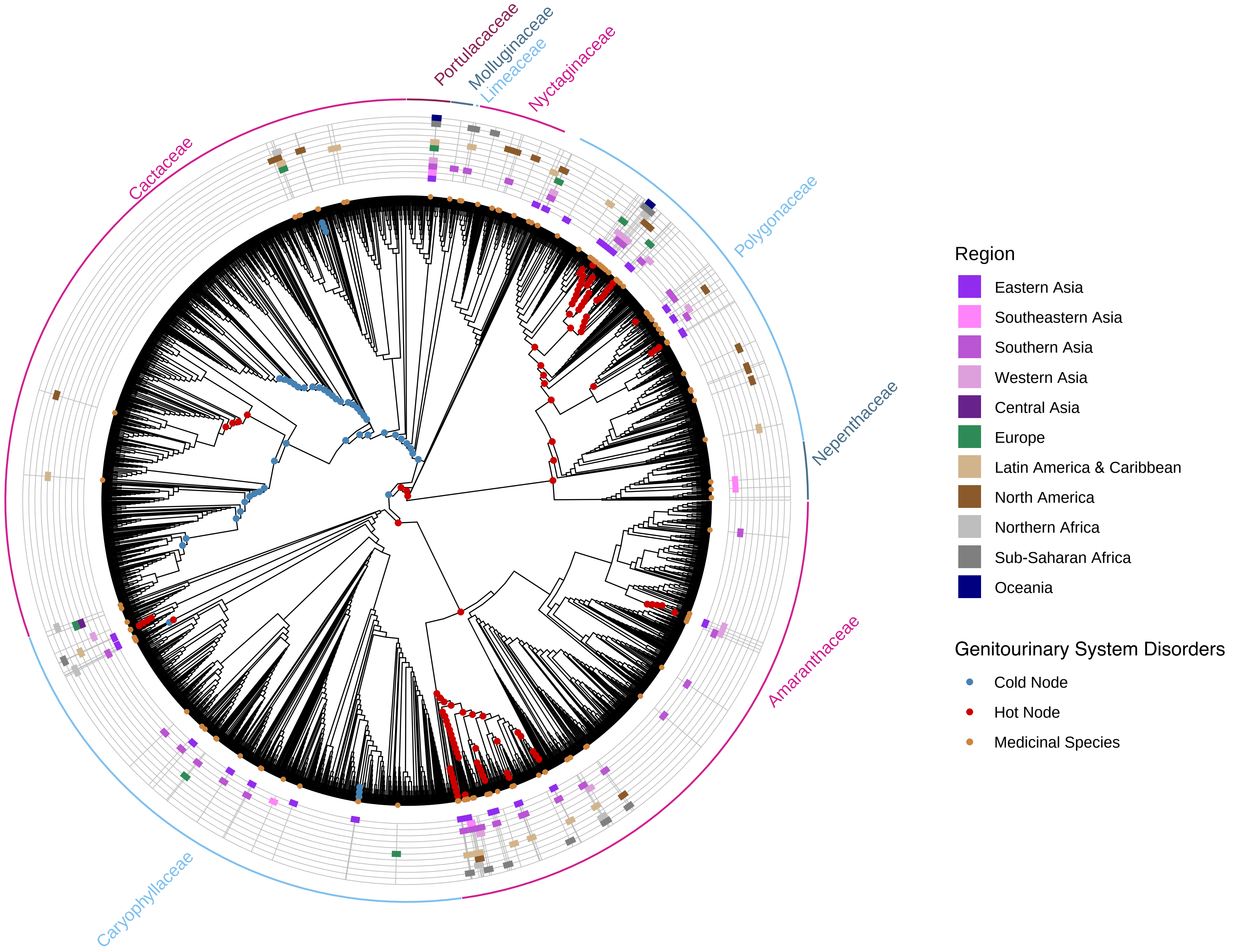
Portulacaceae
Molluginaceae
Limeaceae
Nyctaginaceae
Cactaceae
Polygonaceae
Nepenthaceae
Amaranthaceae
Caryophyllaceae
Region
Eastern Asia
Southeastern Asia
Southern Asia
Western Asia
Central Asia
Europe
Latin America & Caribbean
North America
Northern Africa
Sub-Saharan Africa
Oceania
Genitourinary System Disorders
Cold Node
Hot Node
Medicinal Species

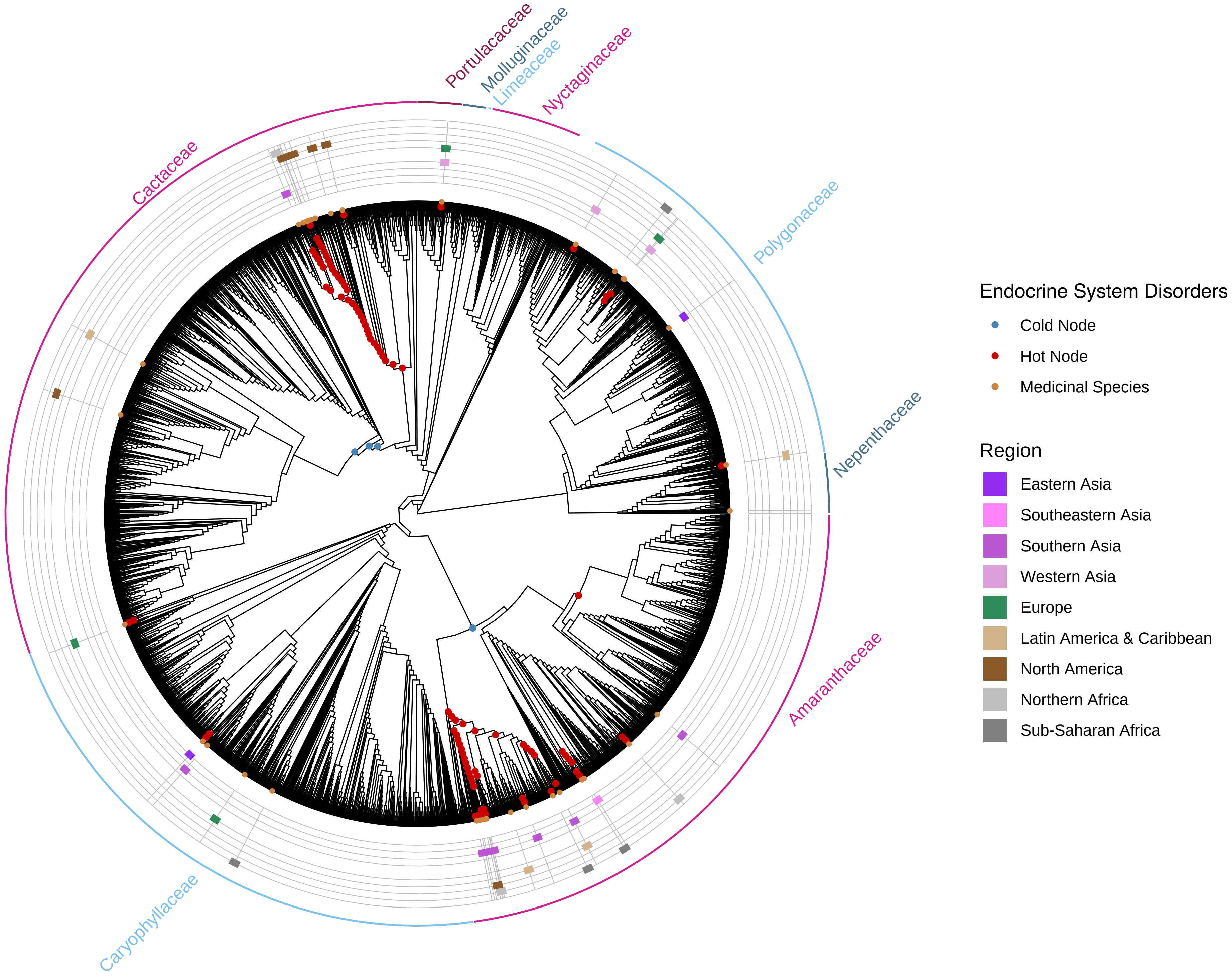

Portulacaceae
Molluginaceae
Limeaceae
Nyctaginaceae
Cactaceae
Polygonaceae
Nepenthaceae
Amaranthaceae
Caryophyllaceae
Endocrine System Disorders
Cold Node
Hot Node
Medicinal Species
Region
Eastern Asia
Southeastern Asia
Southern Asia
Western Asia
Europe
Latin America & Caribbean
North America
Northern Africa
Sub-Saharan Africa

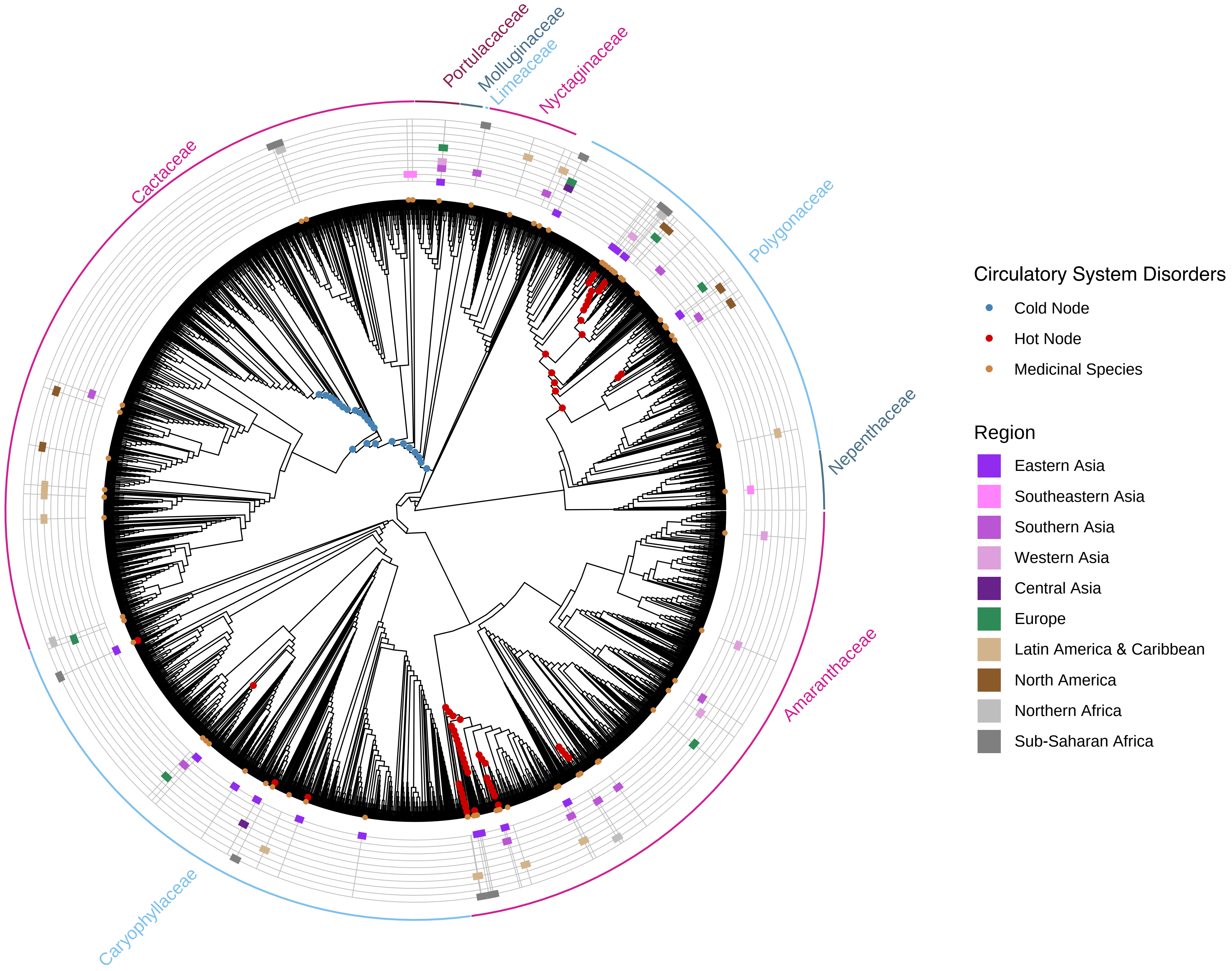

Portulacaceae
Molluginaceae
Limeaceae
Nyctaginaceae
Polygonaceae
Nepenthaceae
Amaranthaceae
Caryophyllaceae
Cactaceae
Circulatory System Disorders
Cold Node
Hot Node
Medicinal Species
Region
Eastern Asia
Southeastern Asia
Southern Asia
Western Asia
Central Asia
Europe
Latin America & Caribbean
North America
Northern Africa
Sub-Saharan Africa

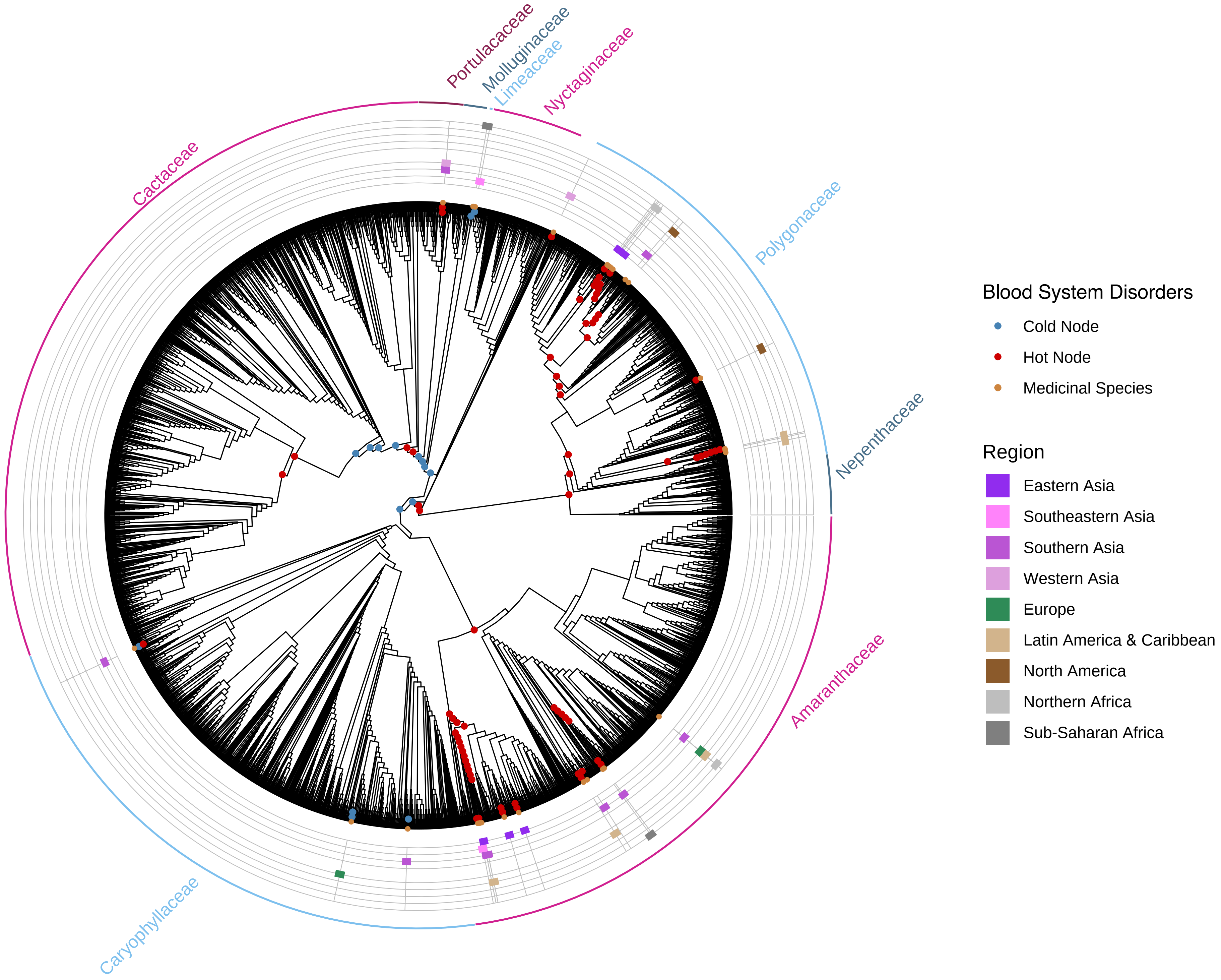

Portulacaceae
Molluginaceae
Limeaceae
Nyctaginaceae
Cactaceae
Polygonaceae
Nepenthaceae
Amaranthaceae
Caryophyllaceae
Blood System Disorders
Cold Node
Hot Node
Medicinal Species
Region
Eastern Asia
Southeastern Asia
Southern Asia
Western Asia
Europe
Latin America & Caribbean
North America
Northern Africa
Sub-Saharan Africa

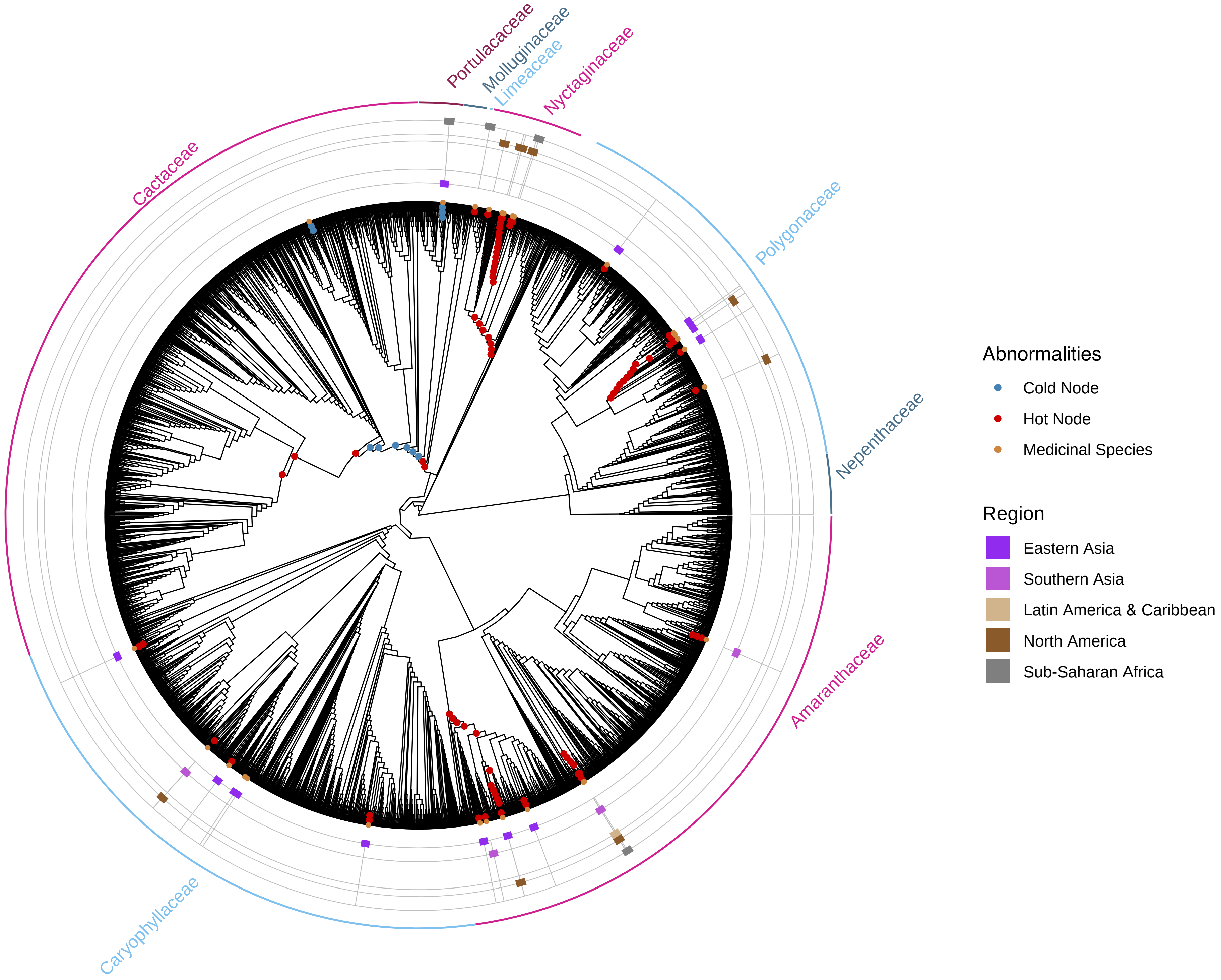

Portulacaceae
Molluginaceae
Limeaceae
Nyctaginaceae
Cactaceae
Polygonaceae
Nepenthaceae
Amaranthaceae
Caryophyllaceae
Abnormalities
Cold Node
Hot Node
Medicinal Species
Region
Eastern Asia
Southern Asia
Latin America & Caribbean
North America
Sub-Saharan Africa
