## Supplementary figures and images for "Traditional medicinal use is linked with apparency, not specialized metabolite profiles in the order Caryophyllales"

### Supplemental 5

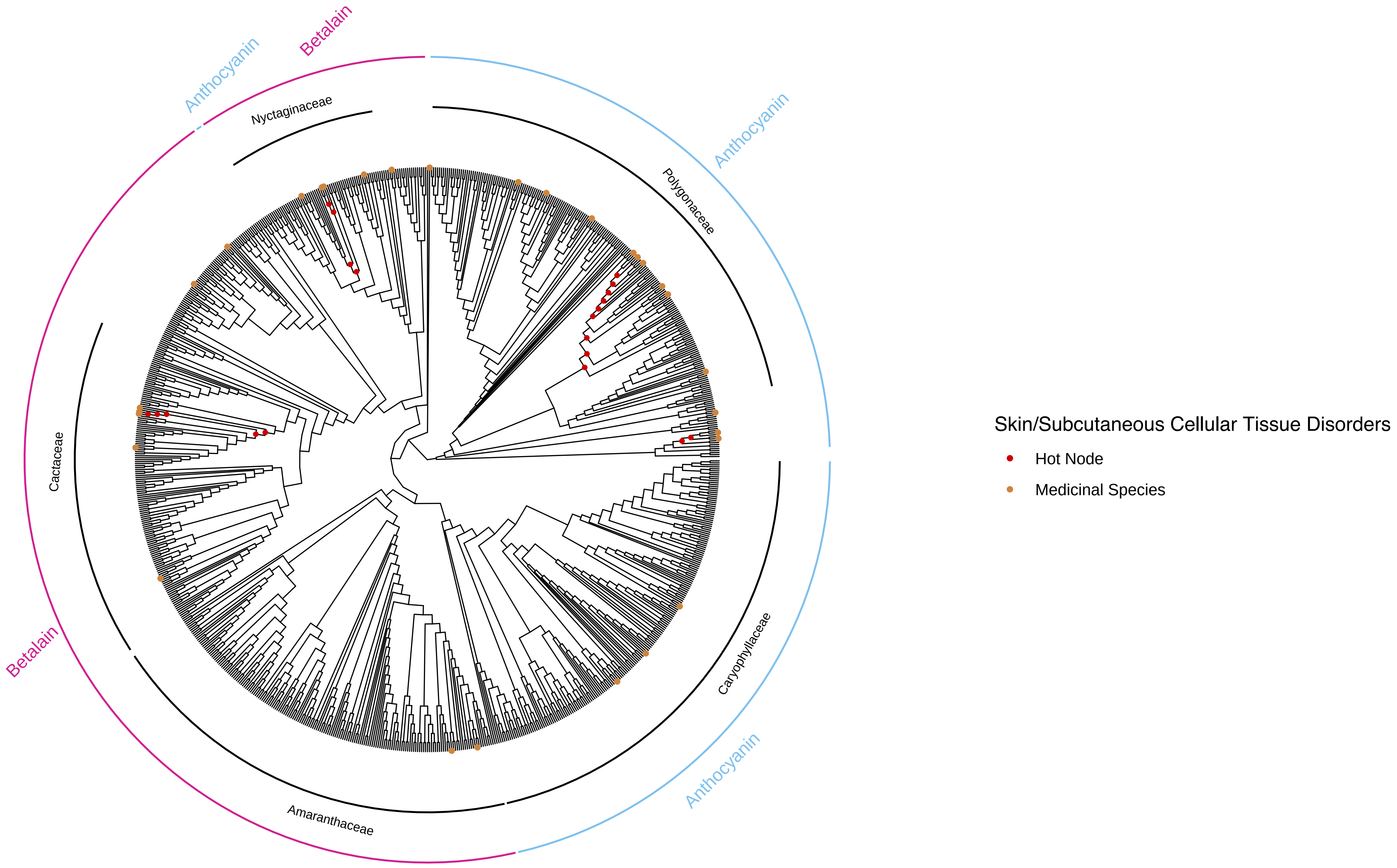

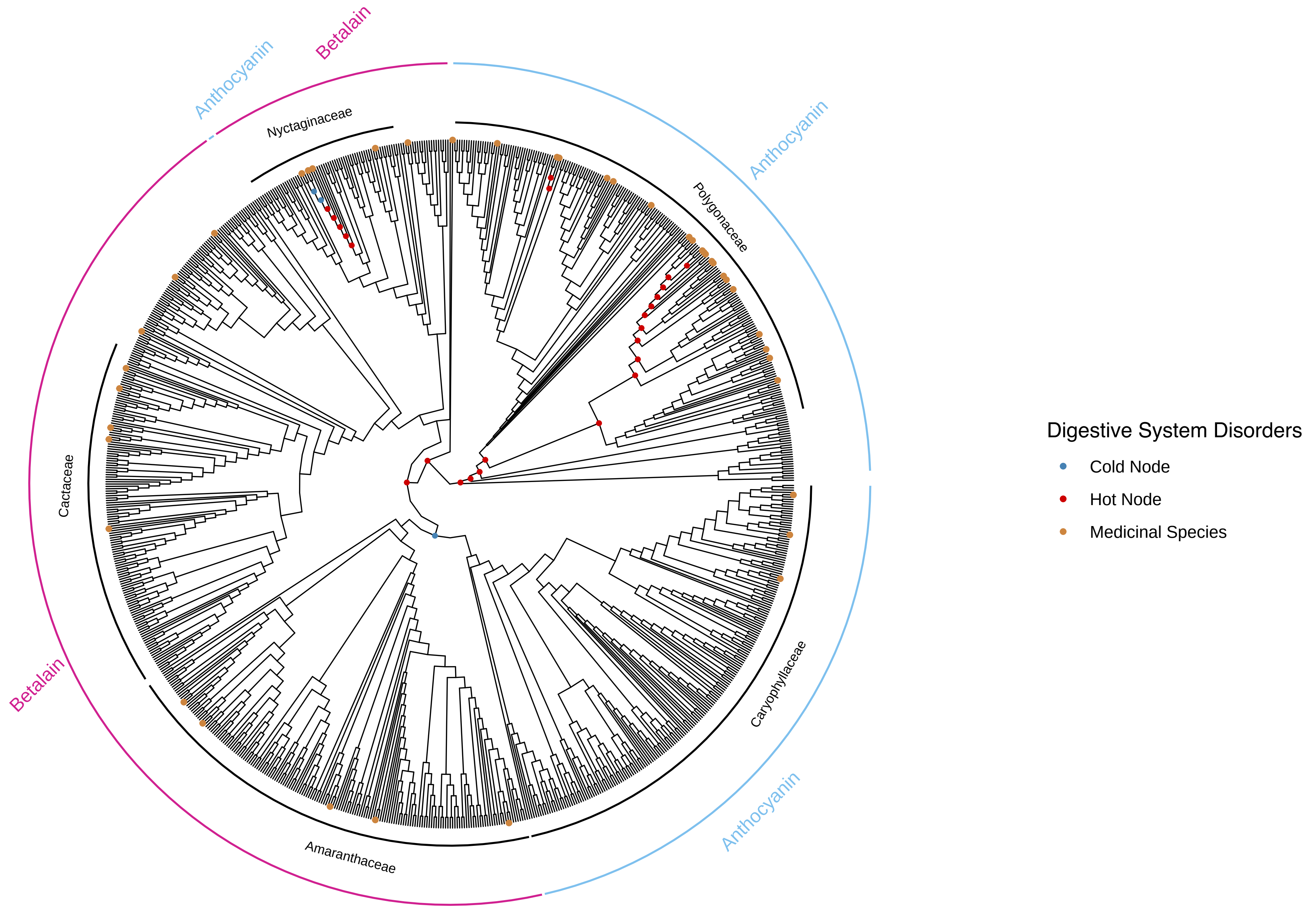

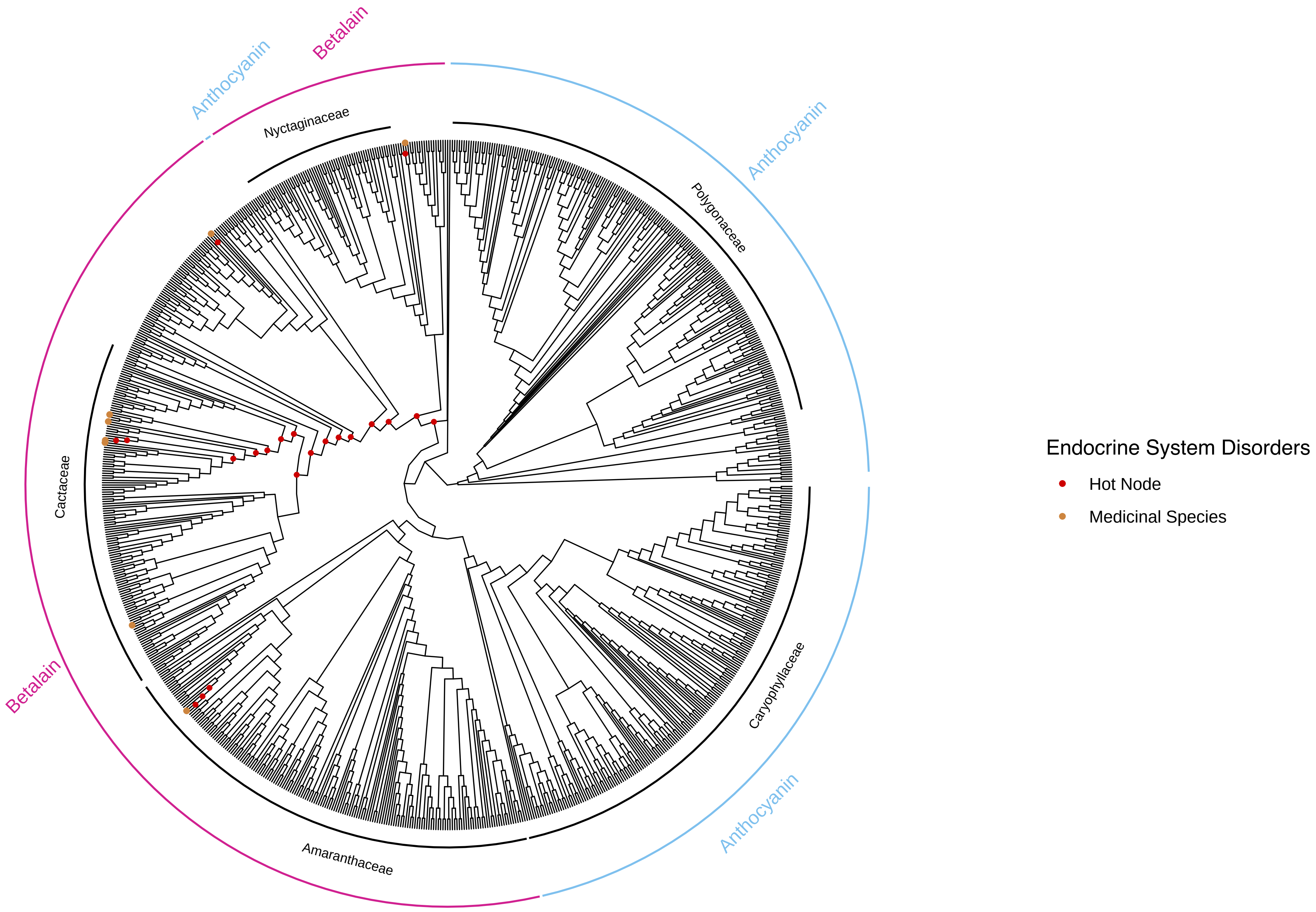

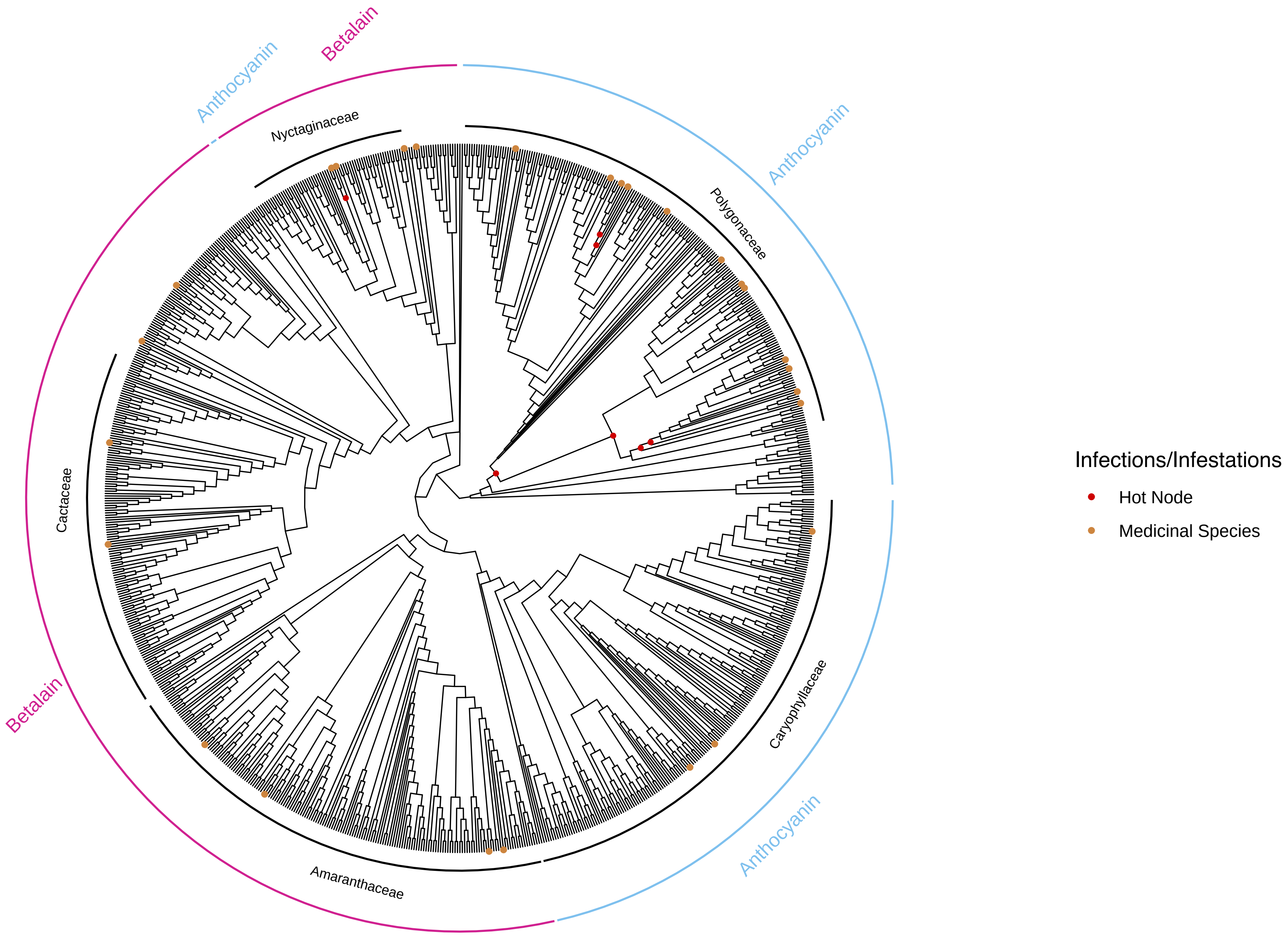

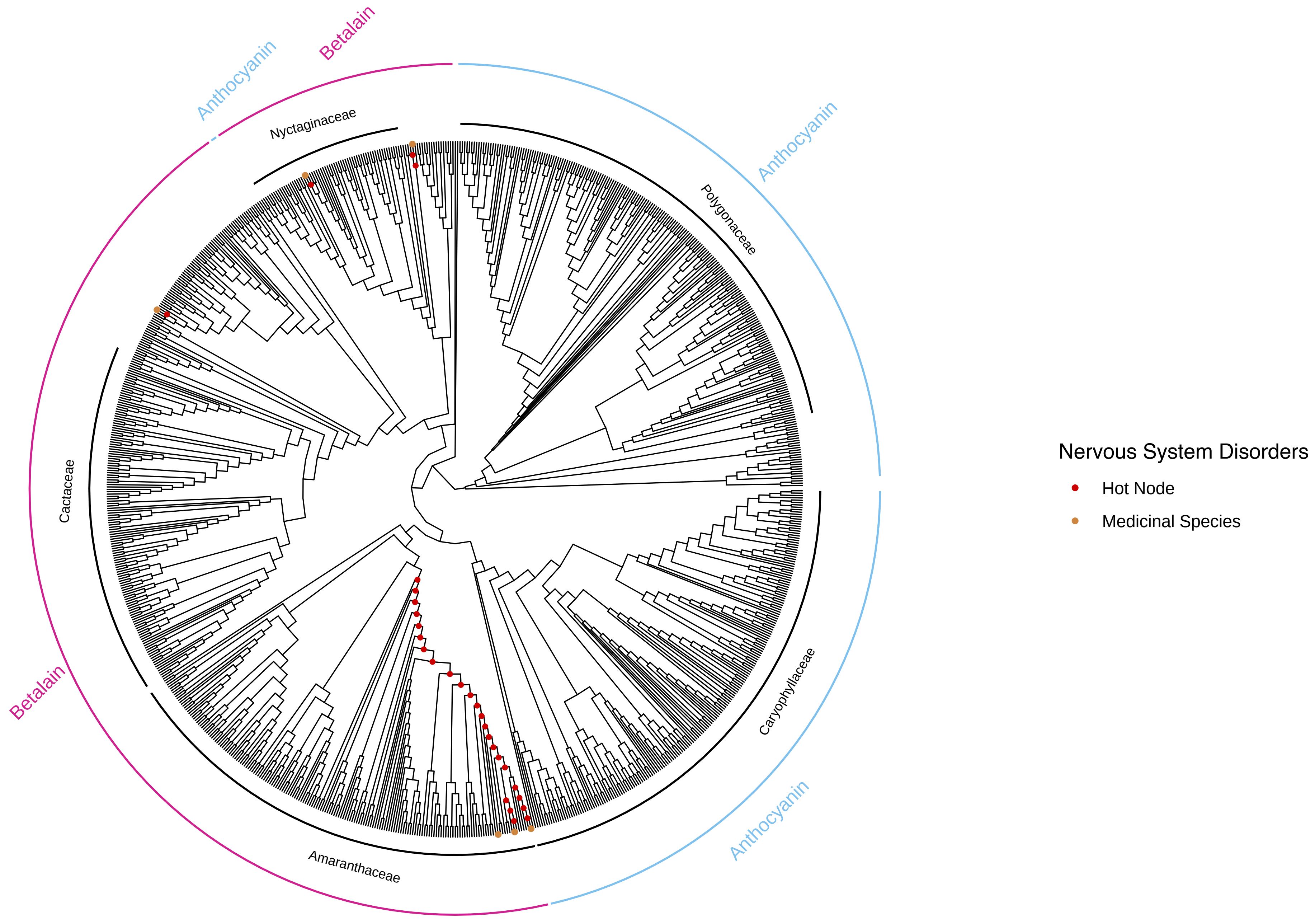

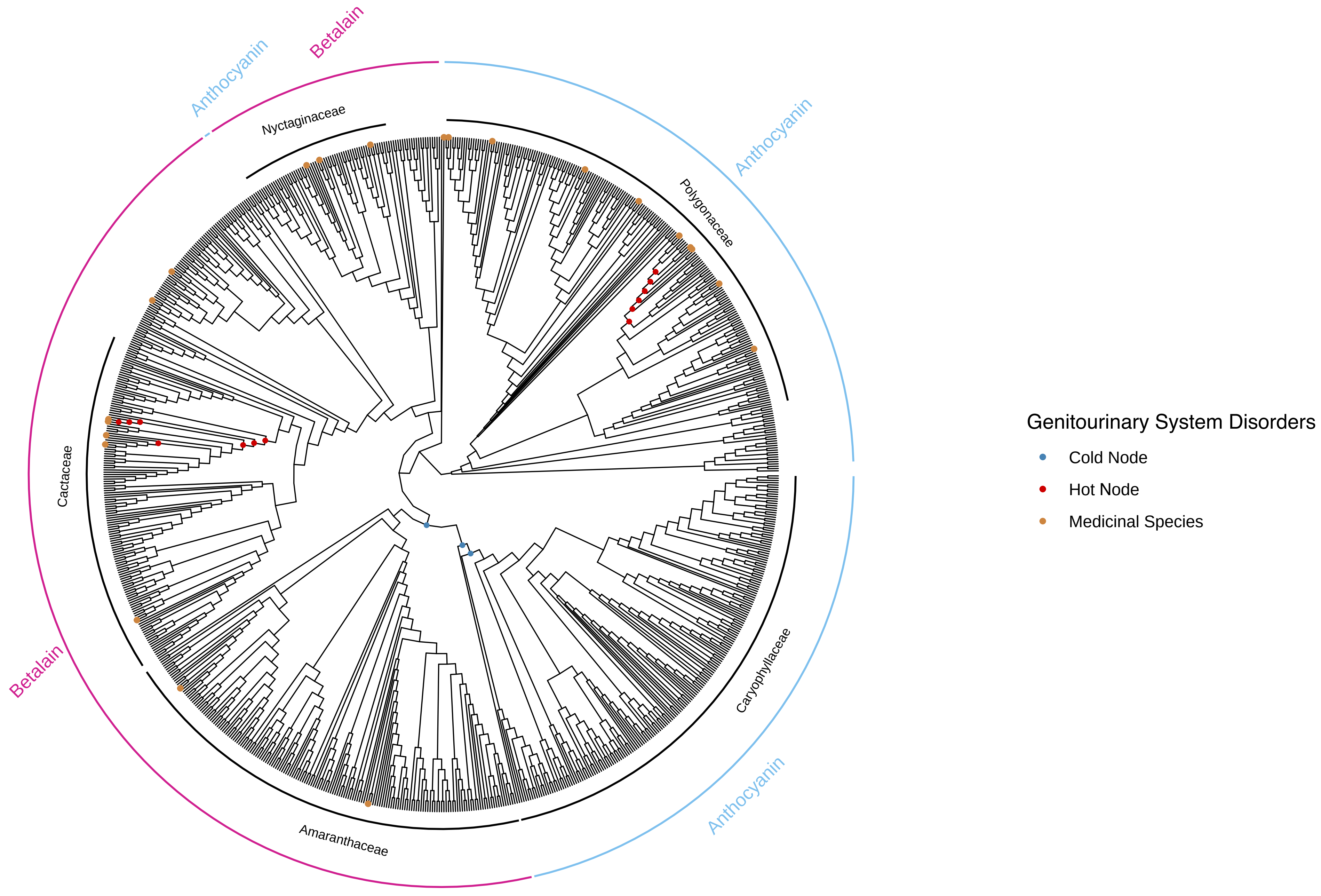

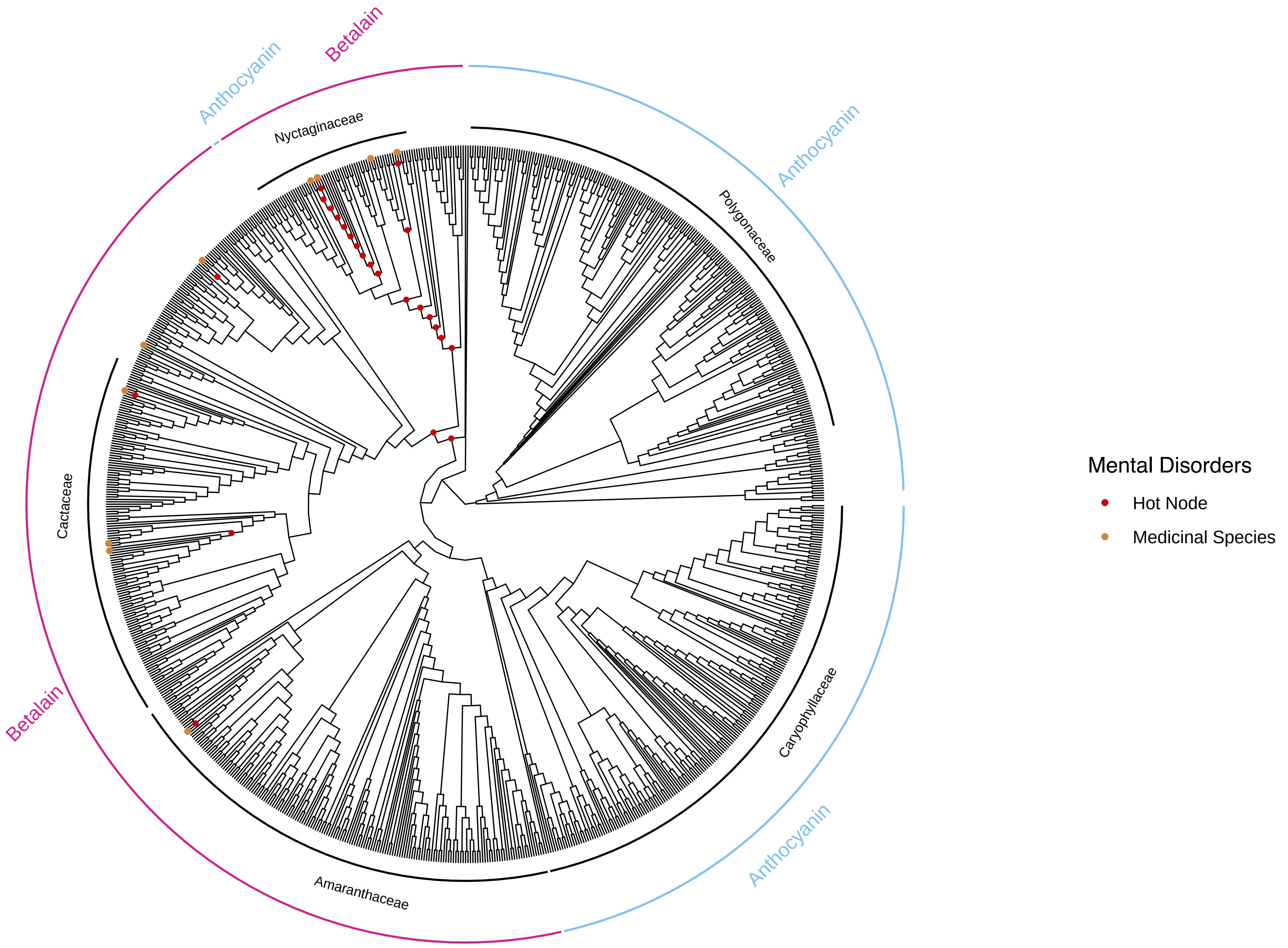

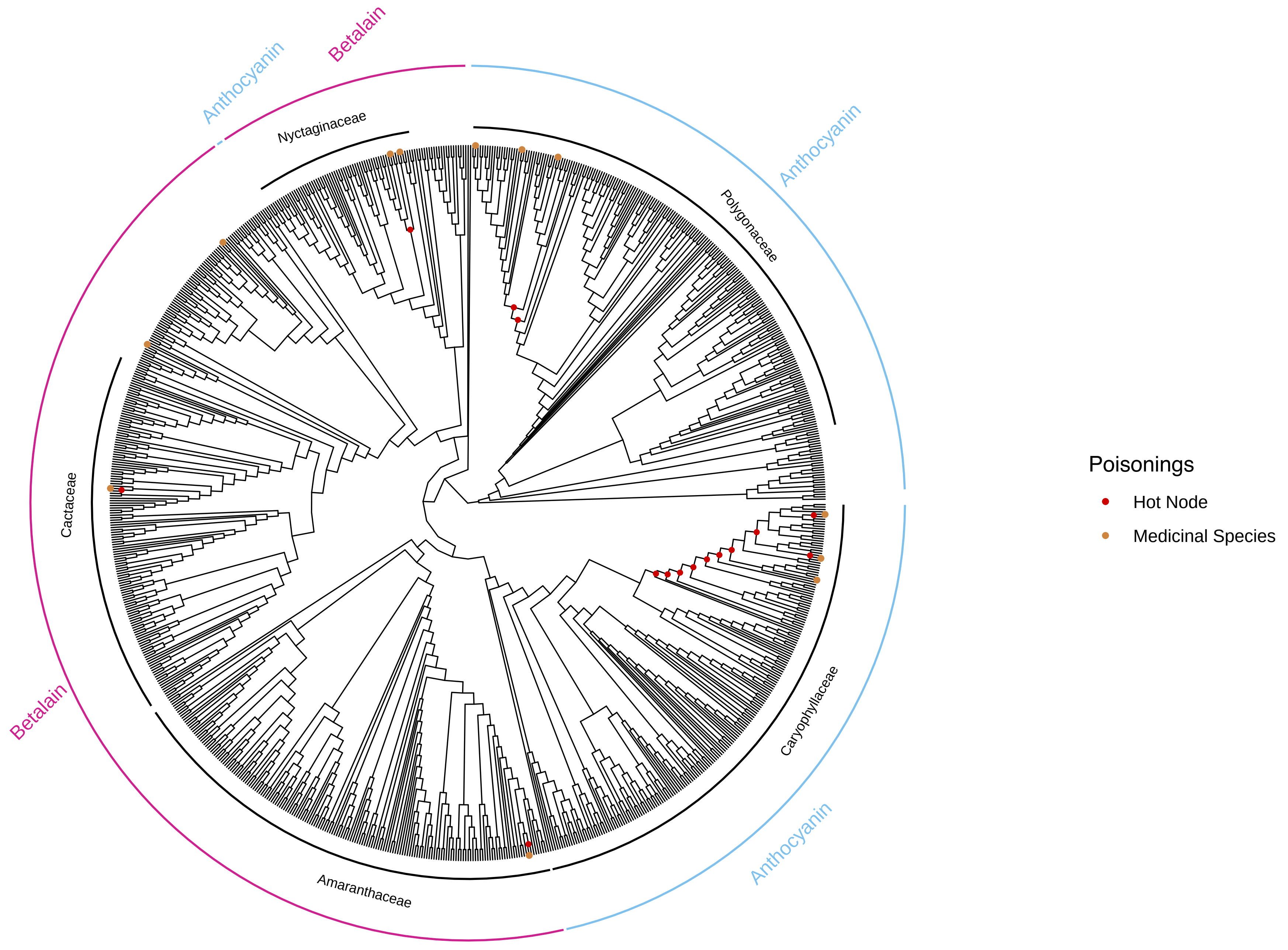

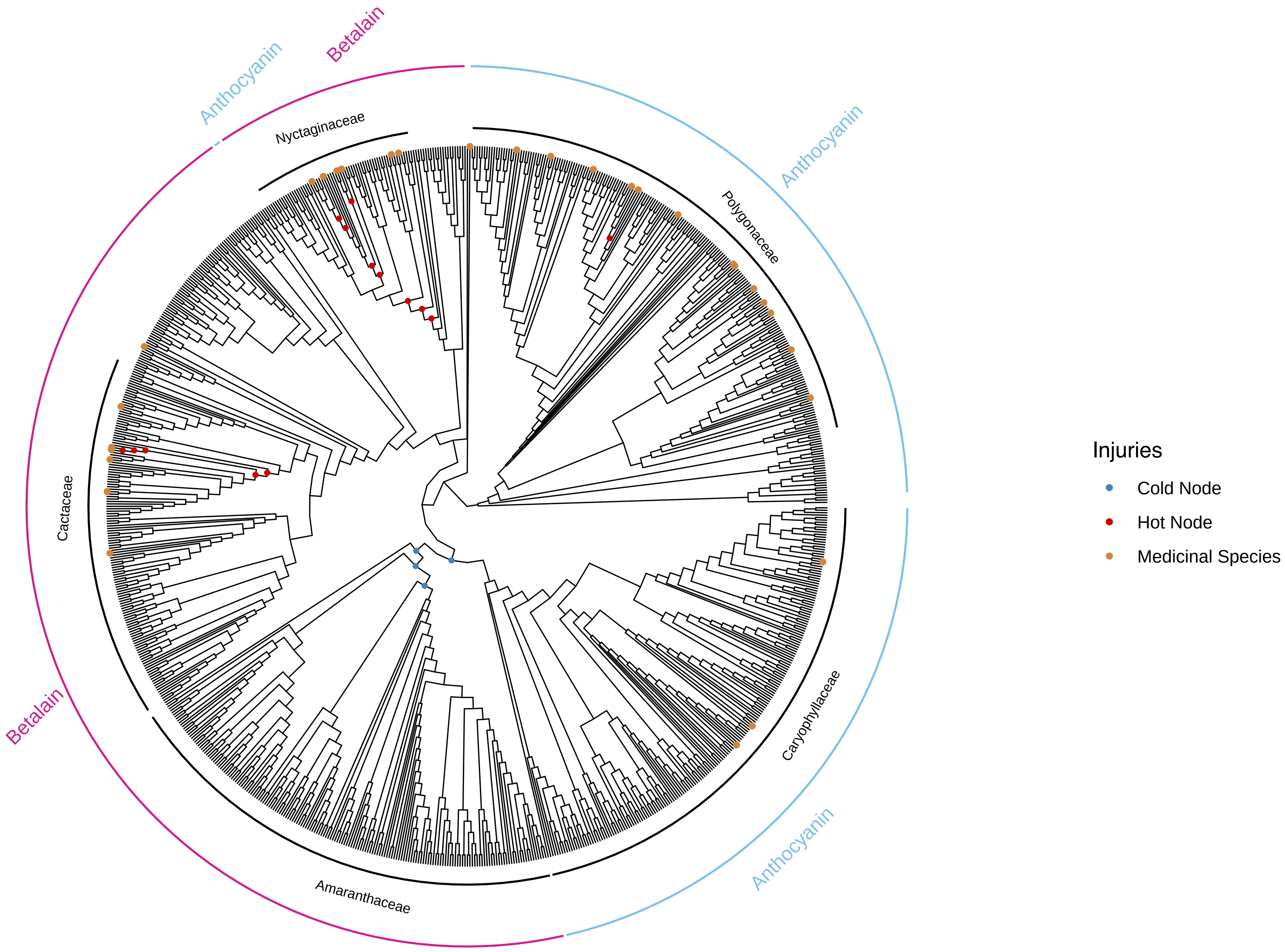
